# A chemigenetic fluorescence lifetime-based biosensor for imaging of ERK activity

**DOI:** 10.64898/2026.10.09.757842

**Authors:** Elias Dressler, Rohan S. Eapen, Dennis Pellanda, Yixin Zhang, Jérôme Julmi, Musadiq A. Bhat, Tommaso Patriarchi, Darren Gilmour, Michelle S. Frei

## Abstract

Extracellular signal-regulated kinase (ERK) governs many cellular processes, and its spatiotemporal activity dynamics are important for downstream signaling specificity. Existing fluorescent ERK activity biosensors, however, fall short in enabling spectrally multiplexed, spatially resolved, or deep tissue measurements. Here, we introduce a chemigenetic, single-fluorophore biosensor, WHaloEKAR, that reports ERK activities through changes in fluorescence lifetime. WHaloEKAR combines large lifetime changes (−0.81 ns) with the spectral versatility of the self-labeling protein HaloTag giving access to orange, red, and far-red ERK biosensors. The latter enabled the investigation of heterogeneous ERK activity at distinct subcellular locations in cultured cell lines and primary cortical rat neurons via fluorescence lifetime imaging microscopy (FLIM). Moreover, multiplexed FLIM measurements allowed to study ERK activity in parallel to the second messengers calcium and cyclic adenosine monophosphate. Finally, WHaloEKAR enabled the investigation of ERK activity in 3D samples as well as in live animals.

## Introduction

Extracellular signal-regulated kinase 1 and 2, collectively called ERK, are key drivers of essential cellular processes such as proliferation, cell survival, and migration. ERK is the final kinase in the RAS-ERK signaling pathway and its activity is tightly regulated via an intricate network of negative feedback loops. ERK activity for instance inhibits the upstream kinases MEK and Raf and phosphorylation of the guanine nucleotide exchange factor SOS prevents effective Ras activation. ERK activity also induces the transcription of dual specific phosphatases that dampen ERK signaling^1^. Dysregulation of ERK activity is associated with developmental and neurodegenerative diseases as well as cancer^2^. On a cellular level, ERK activity is known to be heterogeneous with the frequency, duration, and amplitude of ERK activity playing a crucial role in defining the fate of the cell^3,4^. Moreover the subcellular location of ERK and is translocation to various organelles is known to be key to ensure specific downstream signaling outcomes^2,5^. To capture this heterogeneous behavior, it is therefore important to move beyond traditional static bulk measurements as typically performed by Western Blot. The method of choice in this context is fluorescence microscopy paired with fluorescent ERK activity biosensors, that allow measuring single cell ERK activity dynamics non-invasively with high spatial and temporal resolution. Existing biosensors either make use of a Förster resonance energy transfer (FRET)-design, sandwiching an ERK responsive conformational switch between a FRET-donor and a FRET-acceptor^6–9^, or use the ERK dependent translocation of a fluorescent protein between the cytosol and nucleus^10^. The available FRET-based biosensors show small signal amplitudes and occupy broad spectral regions complicating their use for multiplexed experiments when investigating multiple signaling activities. Kinase translocation reporters (KTRs), on the other hand, use a single spectral channel but require labor-intensive image analysis to distinguish nucleus and cytosol which is often challenging in 3D samples, where variable cytoplasmic volumes and complex cell morphology make accurate segmentation difficult^11^. Moreover, a second nuclear marker often needs to be co-expressed to aid segmentation, again blocking an additional spectral channel. In addition, KTRs only report on a cell’s overall ERK activity and cannot be targeted to distinct subcellular locations. Targetable ratiometric ERK activity biosensors that occupy a single spectral channel orthogonal to existing biosensors are hence highly sought after. Single-fluorophore biosensors with a change in fluorescence lifetime rather than fluorescence intensity could fill exactly this gap as fluorescence lifetime is an inherently ratiometric readout^12,13^. Fluorescence lifetime is independent of measurement parameters like changes in excitation light intensity or sensor bleaching, thus enabling robust quantification. However, only a few dedicated single-fluorophore fluorescence lifetime biosensors are available and many of those are based on short wavelength cyan or green fluorescent proteins and none exist to measure ERK activity^14–21^.

Single-fluorophore chemigenetic biosensors using the self-labeling protein tag HaloTag in combination with far-red synthetic fluorophores have been developed in recent years broadening the spectral pallet of available fluorescent biosensors^19,21–26^. Far-red biosensors benefit multiplexing applications and enable measurements that require high penetration depths, minimal phototoxicity, and high signal-to-background ratios. Recently, first chemigenetic fluorescence lifetime biosensors reporting on NAD^+^ (ref ^21^), Ca^2+^ (ref ^19^), Cdc42^18^, and mechanical force^27^ were reported.

Here, we describe the generation and application of a first single fluorophore chemigenetic ERK activity biosensor with a fluorescence lifetime readout. The spectrally tunable biosensor based on HaloTag can be labeled with synthetic fluorophores covering a broad spectral region, featuring far-red and two-photon compatible fluorophores. We demonstrate its use to measure heterogenous ERK activity dynamics by fluorescence lifetime imaging microscopy (FLIM) both in 2D and 3D cell culture as well as in primary neurons. In addition, multiplexed activity measurements and ERK activity measurements in zebrafish embryos highlight the utility of the biosensor in advanced functional microscopy.

## Results

### Engineering of a chemigenetic fluorescence lifetime-based ERK biosensor

To develop a chemigenetic fluorescence lifetime-based biosensor capable of sensing ERK activity, we utilized circularly permuted (cp)HaloTag[141-147] (Fig. 1a) previously employed to generate far-red chemigenetic biosensors^22,24,26^, including a first intensiometric ERK kinase activity reporter^23^. However, the intensiometric ERK kinase activity reporter mostly showed a change in apparent extinction coefficient connected to the open-close equilibrium of the fluorogenic rhodamine fluorophore used, but no change in fluorescence lifetime. To convert it to a fluorescence lifetime-based biosensor, we exploit a photoinduced electron transfer (PET) quenching mechanism by the electron rich amino acid tryptophan that reduces fluorescence lifetime^26^. Such strategies were previously used to generate Ca^2+^ (ref ^17^), NAD^+^ (ref ^21^) and mechanical force biosensors^27^. Specifically, we combined cpHaloTag[141-147] with the far-red but non-fluorogenic fluorophore JF_669_-HaloTag ligand (JF_669_-HTL, Fig. 1b) and introduced a bipartite ERK specific sensing domain, which consists of an ERK specific substrate^28^ as well as the phospho-amino acid binding domain (PAABD) WW^29^. Biosensor variants were expressed in HeLa cells, labeled with JF_669_-HTL and fluorescence intensity was measured as a proxy for fluorescence lifetime before and after stimulation with human epidermal growth factor (hEGF). Via three rounds of rational engineering testing different tryptophan mutations^30^, domain orientations, and linker variants, we identified a biosensor prototype carrying the Q165W mutation on cpHaloTag showing an intensity change of ΔF/F_0_ = −0.22 ± 0.03 (mean ± SEM, *n* = 83 and 44 cells) and a change in phase fluorescence lifetime of Δτ = −0.195 ± 0.005 ns (*n* = 718 and 720 cells, Fig. 1d and Supplementary Table S1). In contrast to previous chemigenetic and fluorescent protein-based kinase activity reporters^23,31,32^, which followed the substrate-cpHaloTag-PAABD organization, this biosensor followed an inverted PAABD-cpHaloTag-substrate arrangement (Fig. 1c-d).

**Fig. 1.**
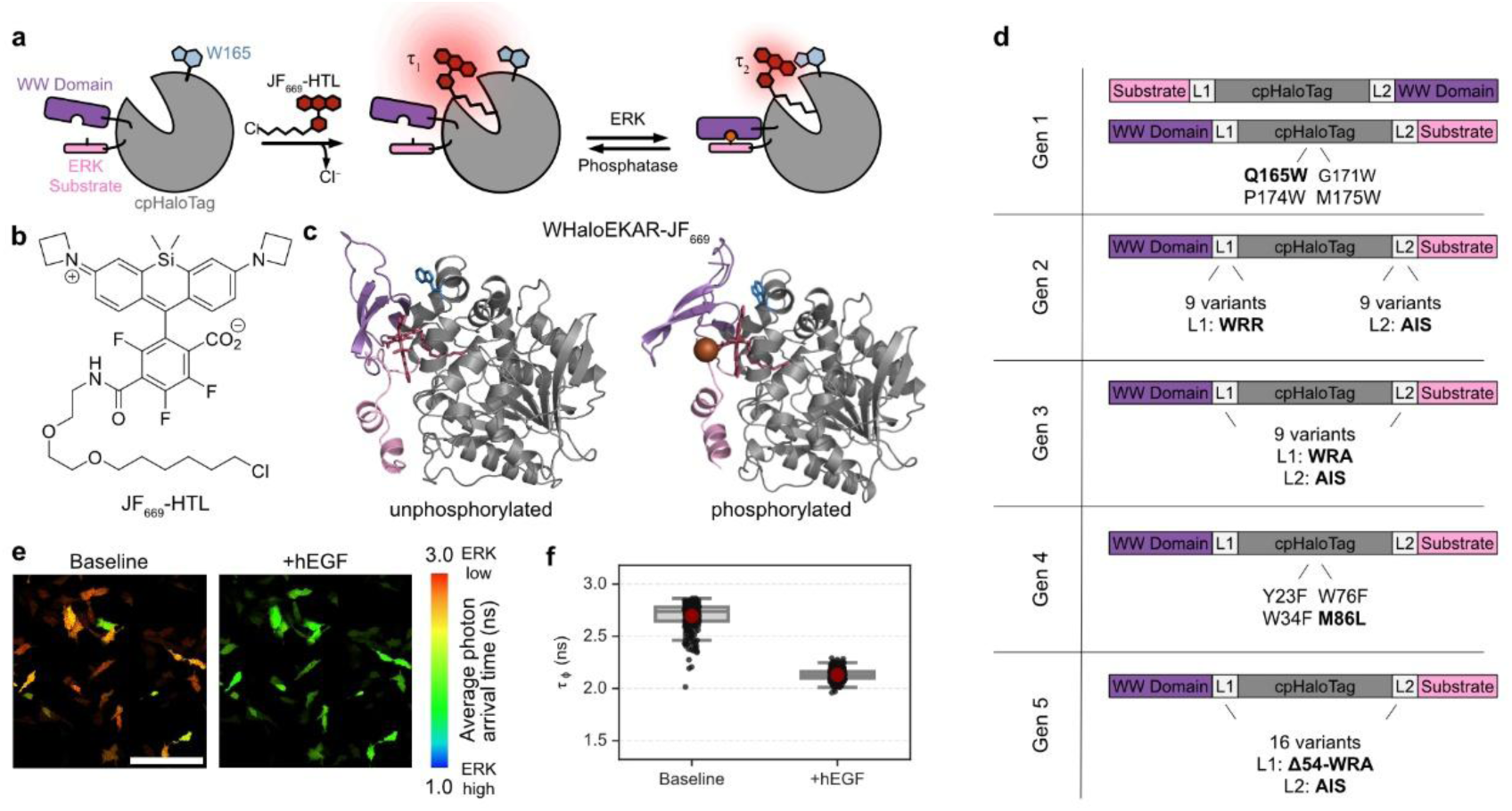
Design and engineering of WHaloEKAR. **a**, Schematic depiction of chemigenetic WHaloEKAR biosensor based on cpHaloTag labeled with JF_669_-HTL and a phosphorylation-dependent switch that influences the fluorescence lifetime of JF_669_. **b**, Chemical structure of JF_669_-HTL. **c**, Model of phosphorylated and unphosphorylated WHaloEKAR-JF_669_. The protein is represented as a cartoon, cpHaloTag in gray, ERK substrate in pink, and the WW domain in purple. The mutated tryptophane is represented as light blue sticks, threonine as pink sticks, the phosphate as a bronze sphere, and JF_669_ as dark red sticks. **d**, Domain structures of the biosensors tested throughout the screening campaign. **e**, Representative FastFLIM image of HeLa cells expressing WHaloEKAR labeled with JF_669_-HTL before and after stimulation with hEGF (100 ng mL^−1^). Scale bar, 200 μm. **f**, Average phase fluorescence lifetime of WHaloEKAR-JF_669_ before and after hEGF stimulation (100 ng mL^−1^, *n* = 290 and 298 cells from three replicates). Representative images of three replicates are shown. Boxplots: Box: 25, 50, 75 percentiles, red dot: mean, whiskers: 1.5 times the interquartile range above and below the 1^st^ and 3^rd^ quartile.

Using structure prediction algorithms, we predicted structures of the unphosphorylated and phosphorylated biosensor. Inspection of the residues surrounding the fluorophore let us identify four amino acids that could negatively impact the fluorescence lifetime of the biosensor via competing PET processes. We therefore introduced Y23F, W76F, or W34F mutations in the WW domain or M184L in cpHaloTag (Fig. 1d)^33^. The latter led to an increase in the change in fluorescence lifetime reaching Δτ = −0.353 ± 0.006 ns (*n* = 337 and 327 cells). Finally, testing of another 16 protein variants with altered interdomain linkers led to the isolation of WHaloEKAR that shows a change in fluorescence lifetime of Δτ = −0.565 ± 0.008 ns (*n* = 290 and 298 cells) upon hEGF stimulation (Fig. 1c, e-f, Supplementary Table S1). This change is 2–5 times higher than typical FLIM-FRET biosensors used in this context^29,34,35^. WHaloEKAR is therefore a promising biosensor for ERK activity studies via FLIM microscopy, and it was used for all subsequent experiments.

### Characterization of WHaloEKAR in mammalian cells and *in vitro*

We further characterized WHaloEKAR’s performance in HeLa cells and compared it to a state-of-the art green ERK-KTR based on mNeonGreen (mNG)^36^. We co-expressed WHaloEKAR and mNG-ERK-KTR in HeLa cells, labeled the cells with JF_669_-HTL and performed confocal imaging in the green spectral region (mNG) as well as confocal FLIM in the far-red region (JF_669_). The changes in fluorescence lifetime of WHaloEKAR followed a similar kinetic profile as the change in cytosol-to-nucleus (C/N) ratio of the mNG-ERK-KTR upon hEGF stimulation (Fig. 2a-b, Supplementary Video S1-2). Subsequent inhibition with the ERK and MEK inhibitors U0126 and SCH772984 revealed that the biosensor response is reversible (Supplementary Fig. S1). When pretreated with the two inhibitors (U0126/ SCH772984), HeLa cells expressing the biosensor showed no decrease in fluorescence lifetime when stimulated with hEGF at a later time point indicating that ERK activity indeed mediates the change in fluorescence lifetime. Similarly, a negative control biosensor lacking the phosphorylation site (T7A) showed no fluorescence lifetime change upon hEGF treatment (Fig. 2c).

**Fig. 2.**
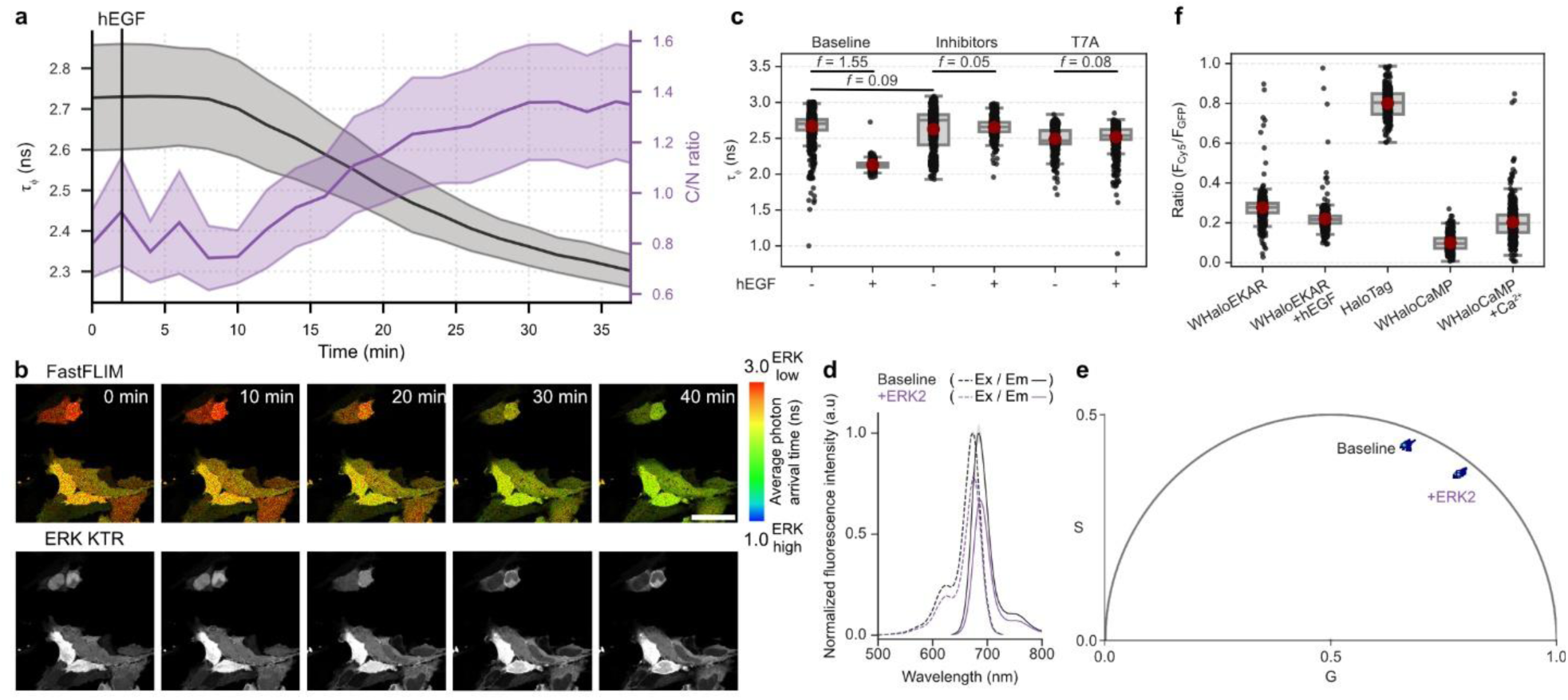
Characterization of WHaloEKAR in HeLa cells and *in vitro*. **a,** Time course of average phase fluorescence lifetime and cytosol-to-nucleus (C/N) ratio of HeLa cells co-expressing WHaloEKAR and mNG-ERK-KTR labeled with JF_669_-HTL upon stimulation of ERK activity with hEGF (100 ng mL^−1^, *n*_φ_ = 9-10 cells, *n*_KTR_ = 11-23 cells). **b,** Representative confocal and FAST-FLIM image. Scale bar, 50 μm. **c,** HeLa cells expressing WHaloEKAR or the dead variant WHaloEKAR(T7A) labeled with JF_669_-HTL upon stimulation with hEGF (100 ng mL^−1^) and inhibition with U0126 (10 μM) and SCH772984 (5 μM, *n* = 1132, 453, 890, 628, 752, and 782 cells). Welch’s one-way ANOVA (unequal variance, two-sided, *F* = 1065.2, DFn = 5, DFd = 1884.2) was performed with Cohen’s f statistics displayed due to large single-cell sample sizes (*t =* −89.24, −3.61, 2.39, 3.16; CI: −0.55–−0.53, −0.06–−0.02, 0.005–0.048, 0.010–0.042; effect size: 1.55, 0.09, 0.05, 0.08, DF = 1525.4, 1435.5, 1269.5, 1521.3; 1–2, 1–3, 3–4, 5–6). **d**, Excitation (dashed) and emission spectra (solid) of purified WHaloEKAR labeled with JF_669_-HTL in the absence and presence of ERK2. **e,** Representative phasor plot of purified WHaloEKAR labeled with JF_669_-HTL in the absence and presence of ERK2. **f,** Brightness comparison of HeLa cells co-expressing EGFP and JF_669_-HTL labeled biosensors or HaloTag7 (*n* = 585, 666, 344, 504, and 567 cells). Time courses are representative of three replicates. Thick solid lines indicate mean response; shaded areas correspond to 95% confidence interval. Representative FastFLIM and phasor images of three replicates are shown. Boxplots: Box: 25, 50, 75 percentiles, red dot: mean, whiskers: 1.5 times the interquartile range above and below the 1^st^ and 3^rd^ quartile, individual data points are shown on top. For *in vitro* spectra mean average of three technical replicates are shown, shaded area indicates SD, as representative data.

Next, we characterized the biosensor *in vitro* using purified WHaloEKAR protein. The biosensor rapidly labeled with JF_669_-HTL reaching an apparent first order rate constant of *k*_app_ = 5.5 ± 0.3 ·10^5^ M^−1^ · s^−1^ (mean ± SD, *n* = 3 measurements), which is only a factor of 10 lower than the literature reported value for HaloTag7 (4.03·10^6^ M^−1^ · s^−1^, Supplementary Fig. S2)^37^. Incubation with active ERK1 or ERK2 led to a change in fluorescence lifetime of −0.81 ± 0.09 ns (mean ± SD, *n* = 4) corresponding to the full dynamic range of WHaloEKAR. We additionally noticed a temperature dependence of the biosensor’s fluorescence lifetime, whereby higher temperatures shift both the non-phosphorylated and phosphorylated states towards shorter lifetimes, while the maximal lifetime change remains largely unaffected. This is in line with previous reports on the fluorescence lifetime of rhodamines at different temperatures and thought to be a result of increased non-radiative deactivation at higher temperatures^38^. The biosensor did not show any changes in absorbance spectra or extinction coefficient. The quantum yield, however, changed from 0.22 in the unphosphorylated state to 0.14 in the phosphorylated state, confirming that stimulation of the biosensor leads to increases in non-radiative decay processes such as PET quenching. The negative control biosensor WHaloEKAR(T7A), on the other hand, showed no changes in extinction coefficient, quantum yield, or fluorescence lifetime upon incubation with ERK1 or ERK2 (Fig. 2d-e, Supplementary Fig. S3, Supplementary Table S2-3). Finally, in its dimmer activated state WHaloEKAR-JF_669_ retains one quarter of the brightness of HaloTag-JF_669_ and both the activated and non-activated biosensor are brighter than WHaloCaMP1a-JF_669_ in its brighter Ca^2+^ bound state (Fig. 2f). This makes WHaloEKAR-JF_669_ particularly well-suited for applications where high brightness is required.

### Spectral versatility of WHaloEKAR

As the spectral properties of a chemigenetic biosensor are only determined at the labeling step, we set out to test the fluorescence lifetime response of WHaloEKAR labeled with a panel of spectrally distinct synthetic fluorophores (Supplementary Fig. S4). Rhodamines with a fluorinated benzoate responded particularly well and allowed shifting the spectral properties into the red (JF_632_-HTL, Δτ = −0.28 ± 0.04 ns) and orange (JF_571_-HTL, Δτ = −0.47 ± 0.05 ns) region (Fig. 3a, b). Other rhodamine-based fluorophores also showed changes in fluorescence lifetime but to a lesser degree (Supplementary Fig. S4). The absolute lifetimes before and after stimulation as well as the change in fluorescence lifetime are determined by the fluorophore and its microenvironment. Especially the distance and orientation to PET donors such as the introduced tryptophan (Q165W) as well as the difference in Gibbs free energy associated with electron transfer as described by the Rehm-Weller formula^39^ are crucial. Hence, both differences in steric and electronic properties of the fluorophore influence PET quenching and thereby the resulting fluorescence lifetime. Fluorination of the benzoate lowers the LUMO compared to non-fluorinated analogues making the fluorophore a better electron acceptor potentially increasing PET efficiency. Modifications such as dealkylation (FSiRh_110_) or fluorination of the xanthene at position 2’, on the other hand, lower the HOMO, which leads to a blue shift and an increase in HOMO-LUMO gap, thereby also impacting PET processes^40^. Interestingly, the choice of the substituent on the aniline nitrogen also impacts the baseline lifetime and the lifetime change, indicating that steric effects must also be considered. In addition, we show that HeLa cells expressing WHaloEKAR labeled with JF_669_-HTL, JF_632_-HTL, or JF_571_-HTL can be imaged via two-photon (2P) excitation using excitation wavelengths of 1240 nm, 1190 nm and 1040 nm, respectively (Fig. 3c). These results demonstrate the utility of WHaloEKAR as a spectrally versatile and 2P excitation compatible ERK activity biosensor.

**Fig. 3.**
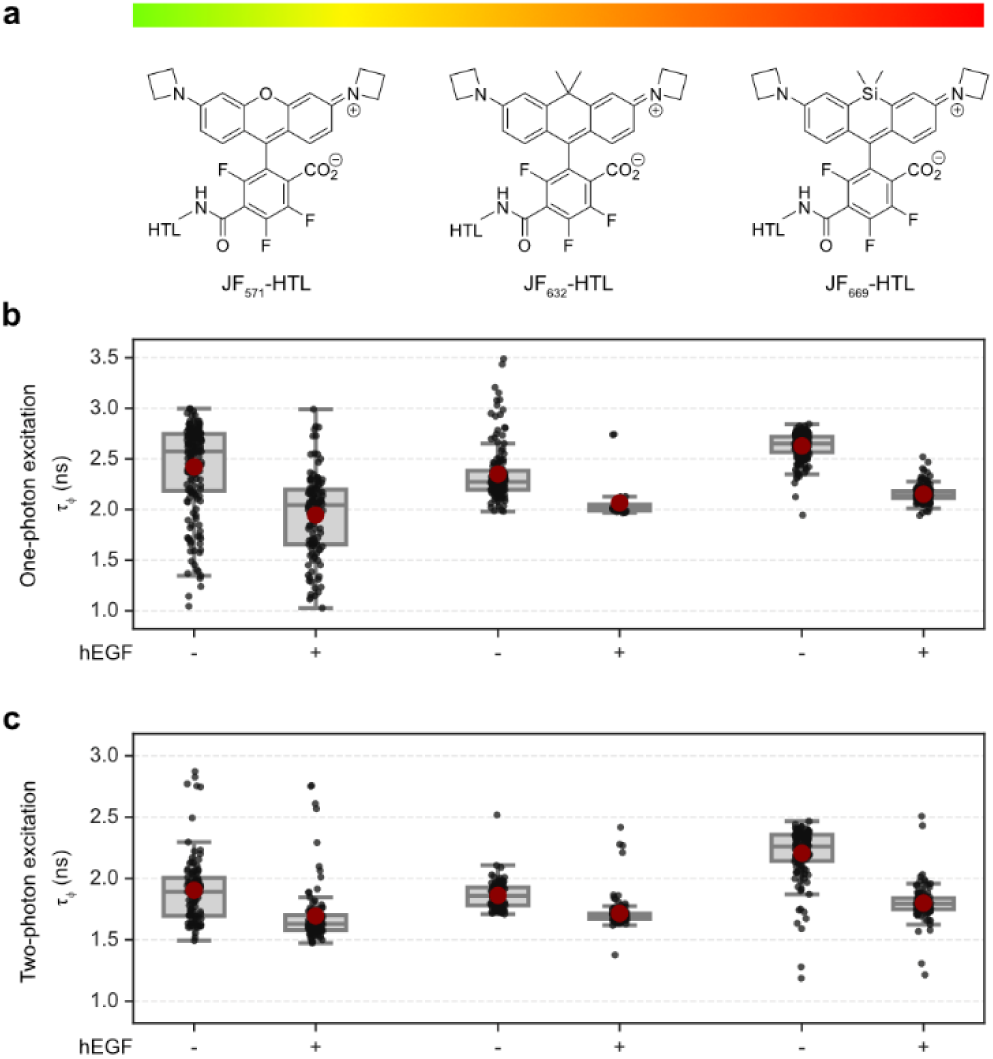
WHaloEKAR is spectrally versatile. **a,** Chemical structures of JF_571_-HTL, JF_632_-HTL, and JF_669_-HTL. **b, c,** Average phase fluorescence lifetime of WHaloEKAR labeled with JF_571_-HTL, JF_632_-HTL or JF_669_-HTL before and after hEGF stimulation (100 ng mL^−1^), measured by one-photon (**b**, *n* = 178, 120, 154, 34, 210 and 208 cells, respectively) or two-photon FLIM microscopy (**c**, *n* = 104, 109, 116, 114, 127 and 110 cells, respectively). Boxplots: Box: 25, 50, 75 percentiles, red dot: mean, whiskers: 1.5 times the interquartile range above and below the 1^st^ and 3^rd^ quartile, individual data points are shown on top.

### Investigation of spatial and temporal ERK activity heterogeneities with WHaloEKAR

To apply our fluorescence lifetime ERK activity biosensor for the investigation of temporal ERK activity fluctuations, we generated a HEK293T cell line stably expressing WHaloEKAR and a plasma-membrane localized EGFP. When labeled with JF_669_-HTL, the bulk of the cells clearly showed an increase in ERK activity upon hEGF stimulation. However, individual cells did not respond (Supplementary Fig. S5, Supplementary Video S3). Upon closer inspection of the unresponsive cell population, they were identified as dividing cells. Indeed, previous reports have shown that hEGF stimulation does not lead to increased ERK activity in mitotic cells due to interference by Cdc2^41^. WHaloEKAR clearly visualized this signaling heterogeneity among different cells. Moreover, ERK activity has been reported to be heterogenous on the subcellular level, with the nucleus, plasma membrane, mitochondria, and various endomembrane compartments showing distinct ERK activity profiles^2,5^. To test if our biosensor could report ERK activity at these distinct locations, we targeted WHaloEKAR to the plasma membrane, lysosomes, endosomes, and outer mitochondrial membrane and labeled it with JF_669_-HTL (Fig. 4a). WHaloEKAR at the plasma membrane, lysosomes, and outer mitochondrial membrane showed lower fluorescence lifetimes when compared to the cytosolic biosensor (NES), indicating higher baseline ERK activity at these compartments (Supplementary Table S4). Inhibition with U0126 (10 μM) and SCH772984 (5 μM) did not lead to statistically significant changes (Fig. 4b, Supplementary Table S4). In contrast, stimulation with hEGF led to a further reduction in mean fluorescence lifetime and hence higher ERK activity at all locations. The change was, however, not statistically significant for mitochondria or the plasma membrane, inferring that under baseline conditions ERK activity almost reached its maximum at the plasma membrane and at mitochondria. Consistent with this interpretation, ERK activity has previously been reported to be higher at the plasma membrane than in the cytosol and to reach near-maximal levels even upon weak pathway stimulation^42^. This behavior has been attributed to scaffolding proteins pre-organizing components of the RAS-ERK signaling cascade as well as to reduced phosphatase activity at the plasma membrane. The latter may also explain why 30 min of pathway inhibition did not result in an increase in fluorescence lifetime as WHaloEKAR reports on the balance between kinase and phosphatase activity. In contrast, less is known about ERK activity at the mitochondria. WHaloEKAR is ideally positioned to provide further insights into ERK signaling dynamics at distinct subcellular locations^2,5^, a measurement that cannot be performed with KTR biosensors. Opposed to FRET-biosensors, WHaloEKAR only occupies one spectral channel, thus opening the possibility to combine multiple biosensors in one and the same or different subcellular location.

**Fig. 4.**
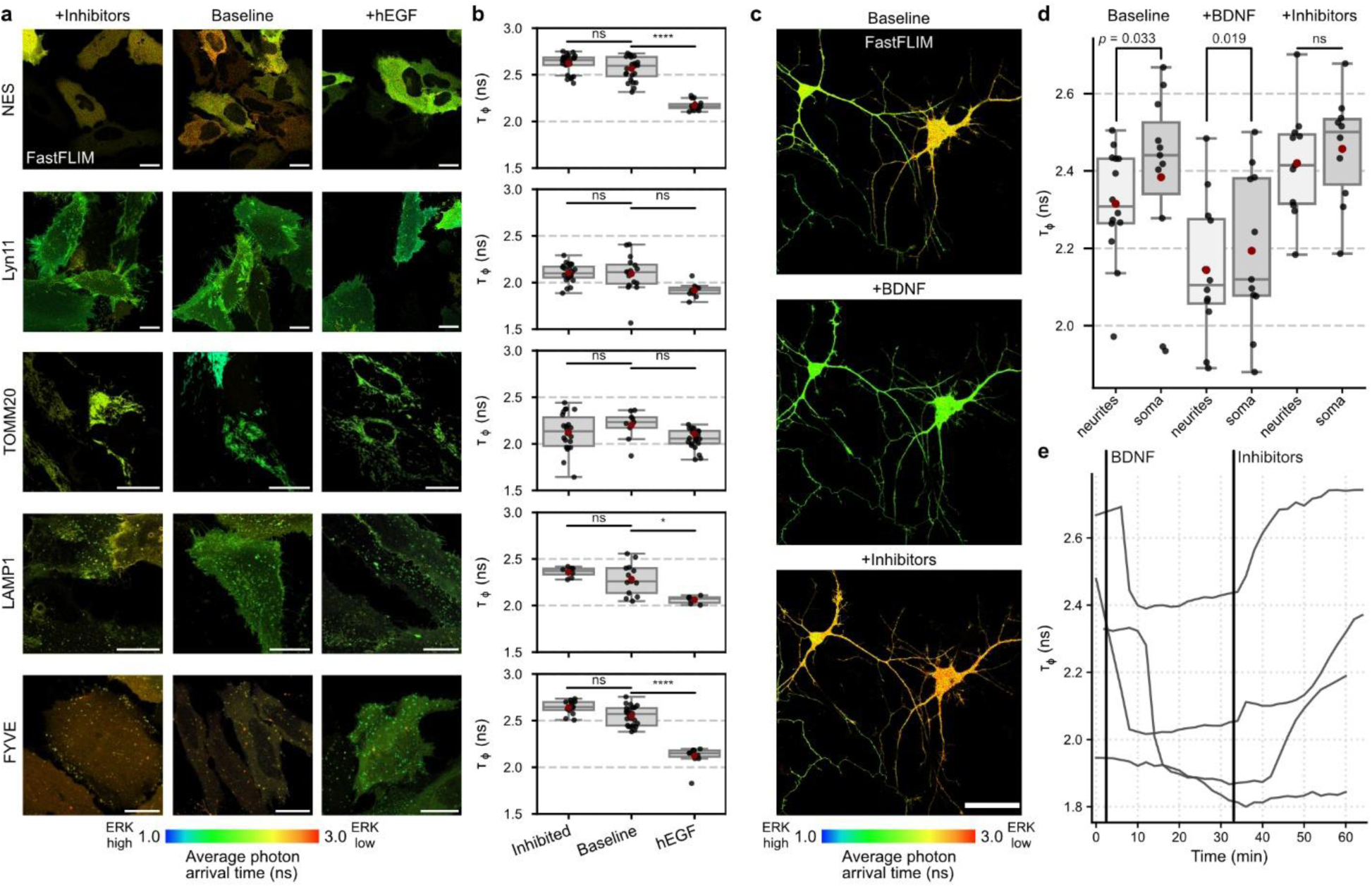
WHaloEKAR reveals subcellular ERK activity heterogeneity. **a,** Representative FastFLIM images of HeLa cells expressing WHaloEKAR at different subcellular locations labeled with JF_669_-HTL. Cells were serum starved (baseline), treated with U0126 (10 μM) and SCH772984 (5 μM) or hEGF (100 ng mL^−1^) for 30 min. Scale bars, 20 μm. **b,** Average phase fluorescence lifetime quantification for WHaloEKAR expressed cytosolic (NES, *n* = 14, 24, 17 cells), at the plasma membrane (Lyn11, *n* = 23, 16, 9 cells), at the mitochondria (TOMM20, *n* = 22, 9, 22 cells), at lysosomes (LAMP1, *n* = 8, 13, 6 cells) and at endosomes (FYVE, *n* = 13, 24, 13 cells). Welch’s one-way ANOVA (unequal variance) with pairwise post-hoc comparisons were used to assess significance. Test statistics are given in Supplementary Table S4. **c,** Representative FastFLIM images of primary cortical neurons expressing WHaloEKAR labeled with JF_669_-HTL at baseline and upon stimulation with brain-derived neurotrophic factor (BDNF, 10 ng mL^-1^) or inhibition with U0126 (2 μM) and SCH772984 (1 μM). Scale bar, 50 μm. **d,** Quantification of subcellular activity heterogeneity comparing activity in the soma and neurites (*n* = 14, 11, 12, 11, 11, 10 neurons). Paired one-sided t-tests were performed (*t =* −2.06, −2.39, −1.22, CI: −0.143–−0.009, −0.079– −0.011, −0.091–0.018, effect size: −0.65, −0.76, −0.41, DF = 10, 10, 9, *p* = 0.033, 0.019, 0.127). **e**, Phase fluorescence lifetime of the soma of four individual neurons upon stimulation with BDNF (10 ng mL^-1^) and subsequent inhibition with U0126 (2 μM) and SCH772984 (1 μM). Representative images of three replicates are shown. Boxplots: Box: 25, 50, 75 percentiles, red dot: mean, whiskers: 1.5 times the interquartile range above and below the 1^st^ and 3^rd^ quartile, individual data points are shown on top. Time courses are shown for four neurons from 3 different neuron-glia co-cultures.

Next, we investigated ERK activity in primary cortical rat neurons in neuron-glia co-cultures. Expression of WHaloEKAR under the human synapsin 1 (hSyn) promotor ensured selective expression in neurons. FLIM measurements of JF_669_-HTL labeled neurons showed clear ERK activation and inhibition upon treatment with brain-derived neurotrophic factor (BDNF, 10 ng mL^-1^) or U0126/SCH772984 (2 μM/1 μM) (Fig. 4c-e, Supplementary Video S4). Interestingly, heterogeneity in ERK activity was visible across different neurons as well as within subcellular regions of single neurons. Specifically, ERK activity in neurites was significantly higher, resulting in a lower lifetime than in the soma across all three tested conditions, baseline, BDNF stimulated, and inhibited (Fig. 4d). These results are in agreement with previous reports on higher abundance of phosphorylated-ERK in neurites over the soma^43^. Taken together, WHaloEKAR is able to resolve temporal and spatial ERK activity heterogeneities among diverse cell lines and primary cells.

### Multiplexed activity imaging by FLIM

WHaloEKAR labeled with JF_669_-HTL is spectrally orthogonal to many fluorescent protein-based biosensors. In particular, combinations with other single-fluorophore fluorescence lifetime-based biosensors would allow using the spectral space more efficiently and enable the measurement of multiple analytes via this inherently ratiometric readout. We therefore sought to investigate whether WHaloEKAR-JF_669_ could be combined with the red Ca^2+^ biosensor RCaMP1h and a cyan cAMP biosensor based on qmTurquoise2 (qmTQ2, Fig. 5a)^16^. We co-transfected HEK293T cells stably expressing WHaloEKAR in the cytosol with the other two biosensors, labeled them with JF_669_-HTL, and imaged them by FLIM in all three channels. Sequential stimulation with hEGF, ionomycin and CaCl_2_, as well as forskolin and IBMX, revealed clear changes in fluorescence lifetime for each biosensor indicating increases in ERK activity, Ca^2+^ levels, and cAMP levels respectively (Fig. 5b, Supplementary Fig. S6a, b). The large spectral separation of the three biosensors makes any bleed-through or spectral crosstalk corrections unnecessary. Hence, other fluorescence lifetime biosensors based on RFPs or mTurquoise2 such as those for Ca^2+^ (ref ^44^) or lactate^45^ could be easily multiplexed with WHaloEKAR-JF_669_. Indeed, co-expressing WHaloEKAR and mTQ2 based Ca-FLITS^44^ in primary rat neurons allowed following changes in Ca^2+^ concentration and ERK activity simultaneously (Fig 5c-d). Depolarization with KCl induced rapid Ca^2+^ transients followed by more gradual increases in ERK activity. The far-red nature of WHaloEKAR labeled with JF_669_-HTL enables extended multiplexing allowing the simultaneous investigation of multiple signaling cascade nodes.

**Fig. 5.**
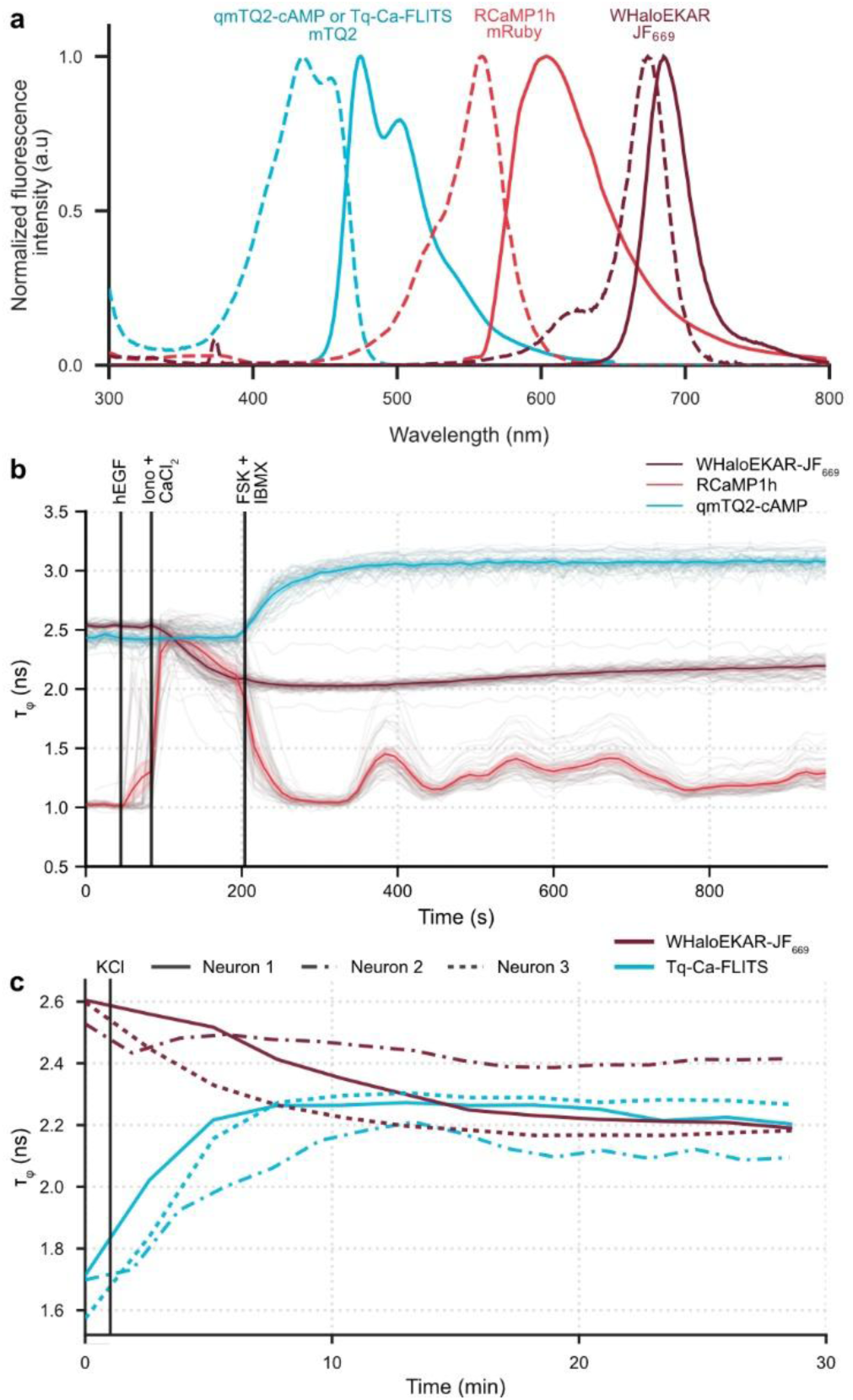
Multiplexed functional microscopy via FLIM. **a**, Excitation and emission spectra of mTurquoise2 (mTQ2), mRuby and JF_669_.^50^ **b**, Three-color multiplexing of cAMP (qmTQ2-cAMP, cyan), Ca^2+^ (RCaMP1h, red) and ERK activity (WHaloEKAR-JF_669_, dark-red) in HEK239T cells stimulated with hEGF (100 ng mL^−1^), ionomycin + CaCl_2_ (1 μM, 5 mM), and FSK + IBMX (50 μM/ 100 μM). All three activities are read out by FLIM (*n* = 64 cells). **c,** Two-color multiplexing of Ca^2+^ (Tq-Ca-FLITS, cyan) and ERK activity (WHaloEKAR-JF_669_, dark-red) in primary cortical neurons stimulated with KCl (50 μM). Both activities are read out by FLIM (*n* = 3 cells). b, Time courses are representative of three replicates. Thick solid lines indicate mean responses; shaded areas correspond to 95% confidence interval; single-cell curves are shown in gray. c, Single curves are shown for individual neurons curves.

### Imaging of ERK activity in 3D and zebrafish embryos

Far-red excitation and emission are ideally suited for imaging in deeper tissue. Additionally, fluorescence lifetime allows circumventing any intensity induced artifacts such as depth dependent scattering or photobleaching. We therefore tested if WHaloEKAR-JF_669_ would enable the measurement of ERK activity within 3D spheroids with average diameters of ∼70 μm. Specifically, 4-day old HEK293T spheroids expressing both WHaloEKAR and EGFP-CAAX were imaged over their entire 3D volume by confocal FLIM (Fig. 6a-c, Supplementary Fig. S7 a, b). Individual cells as identified in the EGFP channel showed clear increases in ERK activity when stimulated with hEGF and could be tracked over 30 minutes. Similarly, measurement of HEK239T spheroids by 2P FLIM clearly revealed ERK activity changes upon hEGF treatment (Supplementary Fig. S7c, d). Next, we measured ERK activity in 6-day old cysts grown from Madin-Darby canine kidney (MDCK) cells^46^ stably expressing WHaloEKAR and EGFP-CAAX. Cysts with diameters between 130–160 μm were labeled with JF_669_-HTL and imaged by FLIM every 2 min for a period of 3 h. Spatial and temporal heterogeneities were identified with individual cells of the polarized epithelial cell layer showing spontaneous changes in ERK activity. This agrees with previous measurements using a cyan/yellow FRET biosensor^47^.

**Fig. 6.**
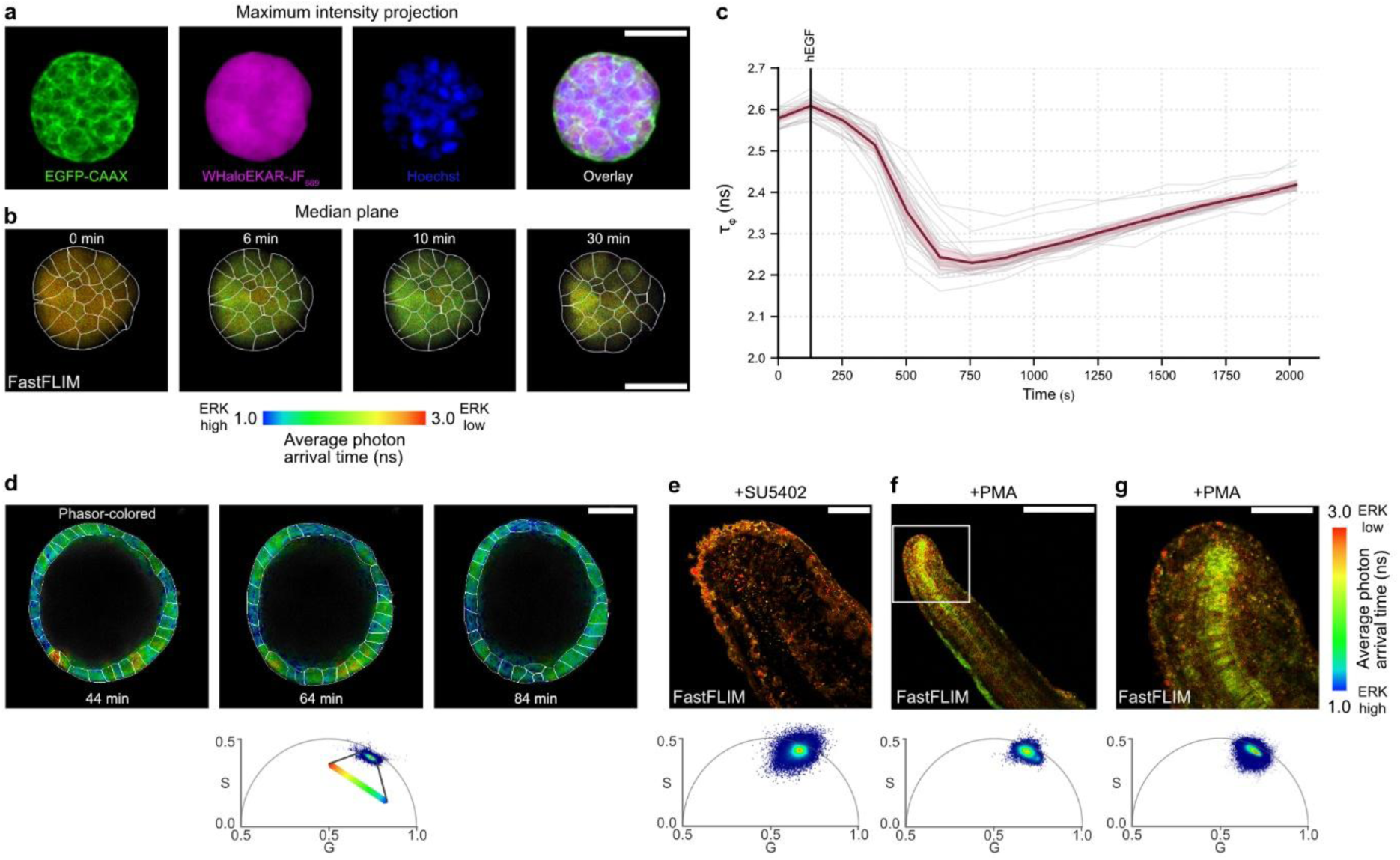
WHaloEKAR reports on ERK activity in 3D spheroids, cysts, and zebrafish embryos. **a, b** Representative confocal maximum intensity projections (**a**) and FastFLIM images (**b**) of 4-day old HEK293T spheroids co-expressing WHaloEKAR and EGFP-CAAX labeled with JF_669_-HTL and Hoechst. Stimulation with hEGF (100 ng mL^−1^) was performed after 2 min. Individual color channels as well as overlay are given. Outline of segmented cells of the median plane shown in **b** are indicated. Scale bars, 40 μm. **c,** Quantification of average phase fluorescence lifetime of all segmented cells within the spheroids shown in (**a**, **b**) over time (*n* = 24 cells). **d,** Representative phasor-colored confocal images of a 6-day old MDCK cyst stably expressing WHaloEKAR and EGFP-CAAX labeled with JF_669_-HTL at three different time-points. Individual cells show spontaneous fluctuations in ERK activity. Phasor plot used to define the look up table is given below. Outline of individual cells as defined by the EGFP channel are given. Scale bars, 40 μm. **e-g,** Representative FastFLIM images of zebrafish expressing WHaloEKAR 24-36 h post fertilization inhibited with SU5402 (**e**) or stimulated with PMA (**f, g**). mRNA was injected at the single-cell stage. The corresponding phasor plots are given. Scale bars, 200 (f) and 50 μm (g). Representative images of three replicates are shown. Time course is representative of three replicates. Thick solid lines indicate mean responses; shaded areas correspond to 95% confidence interval; single-cell curves are shown in gray.

Finally, we investigated the utility of the biosensor for whole animal imaging. We injected WHaloEKAR mRNA into zebrafish embryos at the single-cell stage and measured ERK activity upon indirect activation with the protein kinase C activator phorbol 12-myristate 13-acetate (PMA) that subsequently activates Ras and Raf or after inhibition via the fibroblast growth factor receptor (FGFR) inhibitor SU5402. Imaging WHaloEKAR labeled embryos at 24-36 hours post-fertilization (hpf) revealed distinct patterns of ERK activity when treated with PMA, with cells of the median fin fold as well as notochord cells showing the highest activity levels (lowest lifetime, Fig. 6e-g). In contrast fish pre-treated with SU5402 showed lower and more homogenous ERK activity (Fig. 6e). In combination with recently reported light-sheet implementations of FLIM^48,49^, WHaloEKAR is therefore ideally placed to enable rapid 4D activity sensing of various 3D samples including whole animals.

## Discussion

Despite the central role of ERK activity and in particular its spatial and temporal signaling heterogeneity in governing cellular behaviors such as growth, proliferation, or cell death, its activity is still most commonly measured via bulk methods such as Western blotting or destructive phosphoproteomics. Fluorescent biosensors, on the other hand, enable time-resolved single-cell measurements, but many are currently limited to reporting on cellular but not subcellular activity. Additionally, they use large and blue-shifted spectral regions preventing multiplexed measurements and applications in scattering environments. Here, we present an optimized far-red chemigenetic ERK activity biosensor that allows reading out ERK activity via a robust −0.81 ns *in vitro* change in fluorescence lifetime rather than fluorescence intensity. This inherently ratiometric readout is ideally suited for activity measurements under challenging experimental conditions such as heterogenous light penetration and photobleaching or when comparing samples at distant time points or across laboratories.

Further, WHaloEKAR is spectrally versatile and can be labeled with synthetic fluorophores compatible with both 1P and 2P excitation spanning from 550 to 670 nm. This allows tailoring the spectral properties of the biosensor to individual experiments depending on other biosensors or optical tools used. Accordingly, we show three- and two-color multiplexing of far-red WHaloEKAR with cyan and red, fluorescent protein-based biosensors, reading out all of them by FLIM. This has the potential to revolutionize the study of intricate signaling cascades as multiple nodes of a signaling pathway can be readout simultaneously, providing quantitative information on their abundance or activity helping to inform mathematical modelling of these cascades.

Using WHaloEKAR we show ERK activity heterogeneities on the subcellular level revealing the plasma membrane, lysosomes, and mitochondria as locations of potentially higher ERK activity. Similarly, neurites of primary cortical neurons were found to have higher ERK activity than the neuron’s soma consistent with previous findings. Finally, fluorescence lifetime measurements in the far-red spectral region are ideally suited for functional microscopy in scattering environments where intensity-based strategies traditionally struggle. We show that WHaloEKAR-JF_669_ can be used to measure ERK activity inside of spheroids with diameters of ∼70 μm using both 1P or 2P excitation. Measurements of ERK activity of zebrafish embryos, furthermore, revealed distinct ERK activity states among different cell types upon PMA stimulation. Taken together, these experiments confirm that WHaloEKAR successfully reports on ERK activity dynamics across different cell lines and even organisms as complex as zebrafish.

Even though WHaloEKAR was only engineered for five rounds, it showed a −0.81 ns lifetime change *in vitro*, which translated to an average fluorescence lifetime change upon maximal stimulation by hEGF of −0.57 ns *in cellulo* and overall brightness exceeding the brightness of a similar chemigenetic biosensor for Ca^2+^. Further protein engineering efforts via high-throughput approaches could further enhance the dynamic range, allowing us to measure even more subtle changes in ERK activity. Due to the modularity of kinases activity biosensors, we further anticipate that the biosensor design can be extended to other kinases, in particular to the ones whose substrates are recognized by the WW domain.

To conclude, WHaloEKAR is a powerful chemigenetic fluorescence lifetime-based biosensor enabling the investigation of ERK activity heterogeneities across different length and time scales. It is subcellularly targetable, spectrally versatile, and when used with far-red JF_669_, spectrally orthogonal to many available fluorescent biosensor or optogenetic tools. Together, these properties make the biosensor ideally suited for spatially resolved, multiplexed, and deep tissue experiments and we anticipate the biosensor to help reveal the intricate role of ERK activity across diverse cellular processes.

## Methods

### General considerations

HaloTag fluorophores (JF_669_-HTL, JF_632_-HTL, JF_571_-HTL, FSiRh110-HTL, JF_552_-HTL, JFX_576_-HTL, JF_608_-HTL, JFX_612_-HTL, JF_646_-HTL, JFX_673_-HTL, and JF_698_-HTL)^51,52^ were kindly provided by L. Lavis and Open Chemistry (Janelia Research Campus, HHMI). Fluorophores were prepared as stock solutions in dry DMSO (Sigma) and diluted in the respective buffer such that the final concentration of DMSO did not exceed 1% v/v.

Drug treatments with human epidermal growth factor 100 ng mL^-1^ (hEGF, Qkine Qk011); forskolin 50 μM (Fsk, Adipogen #AG-CN2-0089); 3-isobutyl-1-methylxanthine 100 μM (IBMX, Sigma-Aldrich, #I5879); ionomycin 1 µM or 14 μM (iono, Cayman #10004974); CaCl_2_ 5 mM (Sigma #C4901); U0126 2 μM or 10 μM (Cayman chemical, #70970); SCH772984 1 μM or 5 μM (Cayman chemical, #19166); brain-derived neurotrophic factor 10 ng mL^-1^ (BDNF, PeproTech, # AF-450-02); phorbol 12-myristate 13-acetate 100 ng mL^-1^ (PMA, Adipogen #AG-CN2-0010); SU5402 1 μM (Merck, #SML0443); were performed as indicated.

### Plasmids

A pET51b(+) vector (Novagen) was used for protein production in *E. coli* C41(DE3) (LGC, Biosearch Technologies). Proteins were N-terminally tagged with His_x10_, followed by a TEV cleavage site (ENLYFQ|G).

A pcDNA3.1 vector (Invitrogen) was used for transient expression in mammalian cells. WHaloEKAR and its variants were fused to T2A-EGFP or T2A-EGFP-CAAX, and Lyn11, TOMM20, LAMP1, FYVE, or an NES for localization in mammalian cells. Cloning was performed by Gibson assembly^53^. CAAX (KKKKKSKTKCVIM), Lyn11 (MGCIKSKRKDKDP), NES (MPPLERLTL), and WHaloEKAR(T7A) variants were cloned using overlapping primers. DNA was subsequently transformed into chemically competent DH5α *E. coli* via heat-shock and plated on Lysogenic Broth (LB) agar plates with 100 µg mL^-1^ ampicillin and incubated at 37 °C overnight. Plasmids were purified from overnight cultures in LB medium with 100 µg mL^-1^ ampicillin using ZR Plasmid Miniprep-Classic (Zymo Research).

For stable cell line generation, lentiviral plasmids were cloned via digestion of a pSIN^54^ plasmid with EcoRI and SpeI (NEB) and Gibson assembly^53^ of a PCR-amplified insert.

For expression in neurons biosensors were subcloned via digestion of a pAAV-synapsin plasmid with BamHI and HindIII (NEB) and Gibson assembly^53^ of a PCR-amplified insert.

For expression in zebrafish WHaloEKAR was subcloned into a pCS2+ vector available in the lab. psPAX2 and pMD2.G were gifts from Didier Trono (Addgene plasmid # 12260; http://n2t.net/addgene:12260; RRID:Addgene_12260 and Addgene plasmid # 12259; http://n2t.net/addgene:12259; RRID:Addgene_12259) and directly used for lentiviral production. pcDNA3.1(-)-qmTQ2-cAMP was a gift from Yasushi Okada (Addgene plasmid # 225250; http://n2t.net/addgene:225250; RRID:Addgene_225250)^16^; pCAG cyto-RCaMP1h was a gift from Franck Polleux (Addgene plasmid # 105014; http://n2t.net/addgene:105014; RRID:Addgene_105014)^55^; pCAG-WHaloCaMP1a-EGFP and pAAV-synapsin-WHaloCaMP1a were a gifts from Eric Schreiter (Addgene plasmid # 205313; http://n2t.net/addgene:205313; RRID:Addgene_205313 and Addgene plasmid # 205308; http://n2t.net/addgene:205308; RRID:Addgene_205308)^19^; H2A-mScarletI-P2A-Akt-KTR-mTurquoise2-P2A-ERK-KTR-mNeonGreen and N1-Tq-Ca-FLITS was a gift from Dorus Gadella (Addgene plasmid # 129631; http://n2t.net/addgene:129631; RRID:Addgene_129631^36^ and Addgene plasmid # 191454; http://n2t.net/addgene:191454; RRID:Addgene_191454)^44^; pCDNA5/FRT/TO_HaloTag7_T2A_EGFP, pcDNA5/FRT/TO_LAMP1-HaloTag11-T2A-TOMM20-HaloTag9, and pET51b-His-TEV-HaloTag7 were a gifts from Kai Johnsson (Addgene plasmid # 169325; http://n2t.net/addgene:169325; RRID:Addgene_169325, Addgene plasmid # 175523; http://n2t.net/addgene:175523; RRID:Addgene_175523, ^30^ and Addgene plasmid # 167266; http://n2t.net/addgene:167266; RRID:Addgene_167266)^37^; pcDNA3-endo-EKAR4 and pcDNA3.1-HaloEKAR2.2-T2A-EGFP were gifts from Jin Zhang (Addgene plasmid # 205513; http://n2t.net/addgene:205513; RRID:Addgene_205513^56^ and Addgene plasmid # 216495; http://n2t.net/addgene:216495; RRID:Addgene_216495)^23^ and were either used as entry plasmids or directly for transient transfection.

### Rational engineering

Initial WHaloEKAR constructs were cloned in pcDNA3.1 starting from HaloEKAR2.2 (#216495^31^). Linkers were varied and tryptophan inserted as indicated in Supplementary Table S1. All plasmids were cloned via Gibson cloning. Unless otherwise stated, amino acid numbering relate to the original HaloTag, WW domain and substrate peptide sequences.

### Boltz modeling

The structures of the WHaloEKAR-JF_669_ complex in its phosphorylated and unphosphorylated states were predicted using Boltz-2.^57^ The input consisted of protein sequences and ligand SMILES. The binding of JF_669_ was constrained to pocket residues corresponding to known residues from HaloTag crystal structures,^37^ otherwise the default settings were used.

### Protein production and purification

Proteins were expressed in *E. coli* strain C41(DE3). Cultures were grown in LB medium at 37 °C to an optical density (OD_600nm_) of 0.6, induced by the addition of isopropyl-β-D-thiogalactopyranoside (IPTG, 0.5 mM) and grown at 16 °C overnight. The cells were pelleted by centrifugation (3,100 ×g, 10 min, 4 °C), resuspended in lysis buffer (50 mM Tris-HCl, 300 mM NaCl, 30 mM imidazole, 1 mM TCEP, pH 8.0) containing Complete EDTA-free Protease Inhibitor Cocktail (Roche), and lysed by sonication (5 s sonication, 15 s rest, 8 min total, 15 % power, Branson SFX250) on ice. The cell lysate was cleared by centrifugation (20,000 ×g, 20 min, 4 °C). Clarified lysate was loaded onto HisPur^TM^ Ni-NTA resin (ThermoFisher Scientific) equilibrated with wash buffer (50 mM Tris-HCl, 300 mM NaCl, 30 mM imidazole, 1 mM TCEP, pH 8.0). Unbound protein was washed away using wash buffer and bound proteins were subsequently eluted using a buffer containing 50 mM Tris-HCl, 300 mM NaCl, 500 mM imidazole, 1 mM TCEP pH 8.0. Eluted fractions were analyzed via SDS–PAGE and stained using PageBlue protein staining solution (ThermoFisher Scientific). Elution fractions containing the desired proteins were pooled and then desalted and concentrated, simultaneously using an Amicon Ultra-15 centrifugal filter device (30K MWCO, Millipore) using desalt buffer (25 mM Tris-HCl, 150 mM NaCl, 1 mM TCEP pH 8.0, leading to purity around 90% (verified by SDS-PAGE Coomassie staining). The proteins were flash frozen and stored at −80 °C.

### Fluorescence lifetime measurements (*in-vitro*)

Purified biosensor (5 μM) was incubated with JF_669_-HTL (2.5 μM) in kinase activity buffer (20 mM HEPES, 100 mM NaCl, 10 mM MgCl_2_, 100 μM ATP, 1 mM DTT, 1% (v/v) DMSO, pH 7.4) for 2 hours at 25 °C, shaking at 400 rpm (ThermoMixer C, Eppendorf) occluded from light. Fluorescence lifetime imaging microscopy (FLIM) was performed using a STELLARIS 8 inverted confocal microscope (described below). Samples were excited in a 96-well plate (P96LB-1.5H, Cellvis) at 669 nm and 40 MHz repetition rate (always < 1 photon per laser pulse per pixel acquired using a 20×/0.75 NA dry or HC PL FLUOSTAR 10×/0.3 dry objective (512×512, pixel size = 1137 or 2275 nm, respectively, dwell time = 2.8 μs). Time-correlated single-photon counting data were collected using a HyD X hybrid detector with an emission spectral window set to 682-750 nm. Measurements were performed at ambient room temperature, unless otherwise specified.

For endpoint equilibrium measurements, purified and labeled biosensor (eq. 1 μM JF_669_-HTL) were measured with five frame repetitions, scanning at 400 Hz with a 512×512-pixel grid in the presence or absence of 1 U purified active-ERK2 (DU650, MRC PPU) or 1 U purified active-GST-ERK1 (DU1509, MRC-PPU) in kinase activity buffer. For phosphorylation timecourse experiments, labeled biosensor (eq. 2.5 μM JF_669_-HTL) were added to the imaging dish and the experiment was started, measuring five frame repetitions every 30 seconds for a total of 1 hour. At the four-minute timepoint, the sample was diluted (eq. 1 μM JF_669_-HTL) using 1 U purified active-ERK2 (DU650, MRC PPU) or 1 U purified active-GST-ERK1 (DU1509, MRC-PPU) in kinase activity buffer. All measurements were performed in triplicate.

For temperature dependence experiments, the temperature controllable incubator and stage (Okolab) were set to the desired temperature, and the plate was allowed to equilibrate for 10 minutes before data were collected as described above.

### Excitation and emission spectra

Excitation and emission spectra of labeled biosensors were collected on a multimode plate reader (SPARK®, Tecan) in the presence or absence of 1 U purified active-ERK2 (DU650, MRC PPU) or 1 U purified active-GST-ERK1 (DU1509, MRC-PPU) in kinase activity buffer using the same samples originating from the fluorescence lifetime measurements. Excitation scans were collected in bottom read mode; λ_Em_ 775/20 nm, λ_Ex_ 500-730/10 nm with a 2 nm step, gain 95. Emission scans were collected in bottom read mode; λ_Ex_ 600/20 nm, λ_Em_ 635-800/10 nm with a 2 nm step, gain 95. Excitation and emission spectra were blank-corrected from wells containing kinase activity buffer. All measurements were performed in triplicate.

### Absorbance spectra and Extinction coefficient measurements

Absorbance spectra of labeled biosensors were collected on a multimode plate reader (SPARK®, Tecan) in the presence or absence of 1 U purified active-ERK2 (DU650, MRC PPU) or 1 U purified active-GST-ERK1 (DU1509, MRC-PPU) in kinase activity buffer using the same samples originating from the fluorescence lifetime measurements. Absorbance scans were measured from 500-700/3.5 nm with a 2 nm step. Absorbance spectra were blank-corrected from wells containing kinase activity buffer. All measurements were performed in triplicate. Extinction coefficients were calculated at 669 nm using the Beer-Lambert law (1):

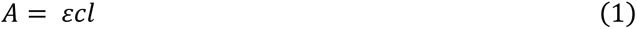

where *A* is the measured Absorbance, *c* is the molar concentration of JF_669_-HTL and *l* is the pathlength in centimeters. The pathlength was calculated based on the volume in a cylinder,

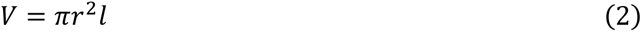

where *V* is the volume of the solution in the well (mL) and *r* is the radius of a single well (0.3 cm, P96LB-1.5H, Cellvis).

### Quantum yield

Quantum yields were determined using a Quantaurus-QY Absolute PL quantum yield spectrophotometer (Hamamatsu). Purified biosensor (5 µM) was incubated with JF_669_-HTL (2.5 µM) in the presence or absence of 5 U purified active-ERK2 (DU650, MRC PPU) or 5 U purified active-GST-ERK1 (DU1509, MRC-PPU) in kinase activity buffer. Samples were measured in a 1×1 cm quartz cuvette excited at 610 nm. Data were analyzed using the in-built software to extract the quantum yields.

### Kinetics by plate reader

Labeling kinetics of WHaloEKAR were measured by time-course fluorescence polarization measurements using JF_669_-HTL. Briefly, fluorescence polarization was measured on a multimode microplate reader (SPARK®, Tecan) in black nonbinding flat-bottom 96-well plates (Greiner) with a final volume of 100 μL at room temperature. Excitation was performed at 620/20 nm, emission was collected at 680/30 nm. The gain and measurement height were optimized from reference measurements (free JF_669_-HTL in labeling buffer (20 mM HEPES, 100 mM NaCl, 0.5 mg mL^-1^ BSA, pH 7.4) and fully bound JF_669_-HTL to WHaloEKAR in labeling buffer). The G-factor was calibrated from the same reference measurements. Kinetic measurements were initiated by injecting WHaloEKAR (final concentration 200 nM) into a well containing JF_669_-HTL (final concentration 50 nM) in labeling buffer, using the built-in injector module. Measurements were performed in triplicate.

The data were fit using a simplified kinetic model (3) in Prism (GraphPad, version 11.0.0) as previously described^37^. Fitting was constrained to minimum and maximum values for the free JF_669_-HTL and the fully bound JF_669_-HTL to WHaloEKAR. *P* and *S* were constrained to concentrations of WHaloEKAR and JF_669_-HTL. A time offset parameter was introduced to account for dead time between injection and the first measurement.

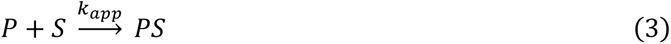

### General cell culture and transfection

HeLa (DMSZ ACC-57), HEK293T (DMSZ ACC-635) or MDCK (NBL-2, ATCC-CCL-34) cells were cultured in Dulbecco’s modified Eagle medium (DMEM, Gibco) containing 4.5 g L^−1^ glucose, 10% (v/v) fetal bovine serum (FBS, Gibco) and 1% (v/v) penicillin-streptomycin (Pen-Strep, Gibco). All cells were maintained in a humidified incubator at 37 °C with a 5% CO_2_ atmosphere. Cells were passaged every 2-4 days or at confluency. Cell lines were regularly tested for mycoplasma contamination via DNA staining.

For imaging experiments, cells were seeded onto sterile quartered 35-mm glass-bottomed dishes (D35C4-20-1.5-N, Cellvis) and grown to 50–70% confluence. For HEK293T cells, dishes were poly-*d*-lysine (ThermoFisher) coated before seeding. Transient transfection of HeLa or HEK239T cells was performed using Polyjet (SignaGen). Cells were changed to fresh serum containing DMEM immediately before transfection. DNA (250 ng) was mixed with serum-free DMEM (12.5 μL), and Polyjet (1 μL) was mixed with serum-free DMEM (12.5 μL). The solutions were then mixed and incubated at RT for 10 min. The transfection complex was added to the wells and incubated for 16 h. Medium was changed to fresh serum-containing DMEM.

Transfection for multiplexing experiments (HEK293T) imaging was also performed using Polyjet (SignaGen) using a total of 250 ng (2 x 125 ng) plasmid DNA.

### Stable cell lines

HEK293T and MDCK cells stably expressing WHaloEKAR-T2A-EGFP-CAAX or WHaloEKAR were generated via lentivirus. First, corresponding pSin plasmids were used to generate lentivirus in HEK293T cells. Briefly, a T25 of 60% confluent HEK293T cells were co-transfected with the pSin plasmid, the viral packaging plasmid psPAX2 (Addgene # 12260) and the envelop plasmid pMD2.G (Addgene # 12259) using Polyjet (SignaGen Laboratories). After 6 h of transfection, the medium was replaced with fresh medium. Viral particles were collected 48 h after transfection and filtered through a 0.45 µm filter (Sarstedt). Lentivirus was then added to HEK293T cells at 50% confluency. 72 h after transduction, medium containing puromycin (2-3 µg mL^-1^, ThermoFisher) was added to select for biosensor-expressing cells. Cells were grown to confluency, and expression was verified by fluorescence microscopy.

### Labeling and sample preparation for live-cell imaging

Cells were labeled with the indicated fluorophores (fluorophore-HTL, 40 nM for 3 h) at 37 °C in corresponding cell culture medium, washed twice with the same medium (1 min each, 37 °C) and changed into serum and phenol red free Leibovitz’s L15 media (L15, Gibco).

For ERK imaging, cells were additionally incubated in serum free L15 for 30 min prior to imaging.

### Widefield microscopy

Images were acquired on a Nikon Ti-2 microscope equipped with a CFI Plan Fluor 40×/1.3 NA oil objective and a SPlanFluor 20×/0.45 air objective, a ORCA-Flash4.0 sCMOS camera (Hamamatsu) and a SOLA light source (Lumencore) controlled by NIS Elements 4.51.01 software (Nikon). Far-red intensity was imaged using an 638/25 excitation filter, a 625 dichroic mirror, and an 670LP emission filter at 300 ms exposure time (far-red filter set); GFP intensity using an 495/20 excitation filter, a 472 nm dichroic mirror, and an 520/35 emission filter exposing for 200 ms (GFP filter set).

#### Time course measurements

Images were acquired with the 40× objective every 15 s for 30 min. Stimulation was performed during the running experiment after 5 min.

#### Brightness comparison with HT7

Far-red and GFP images were acquired with the 20× objective before and after stimulation with hEGF (WHaloEKAR) or ionomycin and CaCl_2_ (WHaloCaMP1a). Raw fluorescence images were first background-corrected by subtracting the fluorescence intensity measured in a cell-free region from the corresponding fluorescence intensity of biosensor-expressing cells in each channel. Brightness was calculated as the ratio of the far-red fluorescence to the GFP fluorescence (Far-red/GFP).

For time-course measurements, the resulting fluorescence values were normalized to the initial fluorescence intensity (F₀), measured at the beginning of the experiment (time zero), and expressed as F/F₀. The maximum fluorescence change after stimulation was calculated as ΔF/F₀ = (Fₘₐₓ − F₀)/F₀, where Fₘₐₓ represents the maximum fluorescence value recorded after stimulation and F₀ represents the fluorescence intensity at time zero.

### General confocal FLIM microscopy

Fluorescence lifetime imaging microscopy (FLIM) were performed using a STELLARIS 8 inverted confocal microscope integrated with the FALCON lifetime imaging module (Leica Microsystems) based on a DMi8 inverted microscope body. The system is equipped with a pulsed White Light Laser (WLL) with a tunable excitation wavelength ranging from 440 to 790 nm; a 405 nm laser diode; HyD X and HyD R photon-counting detectors; HC PL FLUOSTAR 10× 0.3 dry, HC PL APO CS2 20× 0.75 dry, and HC PL APO CS2 63×/1.20 objective. The repetition rate was set to 40 MHz unless otherwise stated. The microscope was equipped with a CO_2_ and temperature controllable incubator and stage (OkoLab). Acquisition of micrographs was performed with the Leica Application Suite X (LAS X) software (v4.8 or v4.9).

### ERK activity measurements

Images were acquired with the 20× objective (512×512 pixels, pixel size = 1.137 um, dwell time = 2.8 μs) at room temperature unless otherwise specified. Exciting at 488 and 669 nm, and collecting at 510-580, 685-800 nm. FLIM measurements were performed exclusively in the far-red channel. For endpoint measurements, images were acquired before and after 30 min stimulation with hEGF. For time-course experiments, images were acquired every 5 min for a total of 30 min. Baseline images were acquired separately before stimulation, and for time-course experiments, stimulation was performed immediately before acquisition of frame 1. Inhibitor experiments were performed after incubation with U0126/ SCH772984 (10 μM/5 μM) for 1 h.

### Different fluorophores

HeLa cells expressing WHaloEKAR were labeled with various fluorophores. Images were acquired with the 20× objective (512×512 pixels, pixel size = 1137 nm, dwell time = 2.8 μs). Excitation and fluorescence collection conditions were as follows: JF_571_-HTL: 571 nm ex, 586-800 nm em; JF_632_-HTL: 632 nm ex, 647-800 nm em; FSiRh110-HTL: labeled cells were exited at 611 nm ex, 626-800 nm em; JF_552_-HTL: 552 nm ex, 567-800 nm em; JFX_576_-HTL: 576 nm ex, 591-800 nm em; JF_608_-HTL: 608 nm ex, 623-800 nm em; JF_646_-HTL: 646 nm ex, 661-800 nm em; JFX_673_-HTL: 673 nm ex, 688-800 nm em; JFX_698_-HTL: 698 nm ex, 713-800 nm em.

### Different subcellular locations

JF_669_-HTL labeled HeLa cells expressing WHaloEKAR at different subcellular locations were preincubated with L15 without serum, hEGF (100 ng mL^-1^) or U0126/ SCH772984 (10 μM/5 μM) for 30 min at 37 °C, and then imaged by confocal FLIM. Images were acquired with the 63× objective (512×512 pixels, pixel size = 362 nm, dwell time = 2.8 μs). Exciting at 488 and 669 nm, and collecting at 510-580, 685-800 nm. Segmentation was performed on the EGFP channel.

### General FLIM image analysis

Raw time-resolved photon histograms are converted to phasor coordinates via discrete Fourier transform and are denoised using wavelet filtering^58^. To determine the fluorescence lifetime of individual cells, images were segmented using Cellpose-SAM^59^. Pixels meeting a defined photon threshold are photon-weighted and the resulting average phase and modulation lifetimes per cell are calculated from the G and S coordinates. LapTrack^60^ was used for tracking single cells over a time course experiment. Unless otherwise stated, FastFLIM images (average photon arrival times per pixel) are shown (LAS X Leica).

*In vitro* analysis: Photon counting histograms were fit to obtain intensity-weighted fluorescence lifetimes. These were calculated in the LAS X software (Leica Microsystems) by fitting three components to the *N*-exponential re-convolution model.

### KTR experiments and analysis

HeLa cells transiently expressing WHaloEKAR-T2A-ERK-KTR-mNG were imaged using the 63× objective (512×512 pixels, pixel size = 482 nm, dwell time = 2.8 μs) sequentially exciting at 404, 488, and 669 nm, and collecting at 425-470, 510-580, 680-780 nm at 22 °C or 37 °C. Acquiring FLIM only in the far-red channel. Images were acquired every 2 min for a total of 30 min and cells were stimulated with hEGF at t = 3 min or inhibited with U0126 and SCH772984 after prior hEGF stimulation.

#### Image analysis

FLIM data was analyzed as described above. The cytosol/nucleus ratio was calculated for the green channel. Briefly, both the Hoechst (nuclei) and green channel (entire cell) were segmented using Cellpose-SAM^59^. The nuclei masks were subtracted from the cell masks providing masks of the cytosol only. The cytosol and nuclei masks were matched. Cells without a nucleus or non-transfected cells (nuclear signal but no cytosol signal) were discarded. The remaining masks were used to calculate per cell cytosol/nucleus ratios from the green channel.

### Biosensor multiplexing

Biosensor multiplexing was performed on the STELLARIS 8 FALCON microscope, acquiring FLIM images in three separate sequences every 12 seconds for up to 30 min at 37 °C. Cells were starved for 30 min in Ca^2+^ and Mg^2+^ free HBSS (Gibco) and then imaged using the 63× objective (512×512 pixels, pixel size = 361 nm, dwell time = 2.8 μs) sequentially exciting at 440, 570, and 669 nm, and collecting at 460-530, 580-650, 690-760 nm while acquiring FLIM in every channel. Stimulation was performed after 36 s (hEGF), 84 s (iono 1 µM, CaCl_2_) and 204 s (Fsk, IMBX).

### Spheroids and cyst preparation

HEK293T or MDCK cells stably expressing WHaloEKAR-T2A-EGFP-CAAX were grown into spheroids or cysts over the course of 4 or 6 days, respectively. 35-mm or quartered 35-mm glass-bottom dishes (Cellvis) were coated with a thin layer of Matrigel (Corning), then cured at 37 °C for 30 min. Cells were trypsinized and seeded on top of the cured Matrigel at a density of 1500-2000 cells per quarter or 6000 cell per dish, then grown in DMEM (Gibco) containing 4.5 g L^−1^ glucose, 10% (v/v) FBS (Sigma) and 1% (v/v) Pen-Strep (Sigma-Aldrich) supplemented with 2% Matrigel for 4 or 6 days.

Spheroids or cysts were labeled with JF_669_-HTL (100 nM for 2 h) and Hoechst in the case of spheroids (5 μg mL^−1^ for 2 h) at 37 °C in cell culture medium with 2% Matrigel, washed with cell culture medium, and placed in L15 supplemented with 2% Matrigel (+ 10% FBS for cyst) for imaging.

### Confocal FLIM microscopy of spheroids

Spheroids were imaged at 37 °C using the 10× objective (512×512 pixels, pixel size = 207 nm, dwell time = 2.8 μs) sequentially exciting at 405, 488, and 669 nm, and collecting at 425-470, 510-580, 690-760 nm while acquiring FLIM only in the far-red channel. z-stack images (16 slices) were collected at intervals of 4.2 µm throughout at total depth of 68.5 μm. Z-stacks were acquired every 2 min for a total of 30 min. Spheroids were stimulated with hEGF at t = 2 min. Cells were segmented in each plane using the green channel (EGFP-CAAX).

### 2P FLIM microscopy

2P FLIM was performed using a STELLARIS 8 DIVE upright confocal microscope integrated with the FALCON lifetime imaging module (Leica Microsystems) based on a DM6 CSF microscope body. The system is equipped with a pulsed White Light Laser (WLL) with a tunable excitation wavelength ranging from 440 to 790 nm; a Discovery NX MP laser (680-1300 nm, with a fixed 1040 nm line) and a Chamelion vision II (680-1080 nm), HyD RLD photon-counting detectors; and a 25×/0.95 NA WATER HC FLUOTAR VISIR 0.17 objective. 2D HeLa cell images were acquired at 512×512 pixels, pixel size = 867 nm, dwell time = 2.8 μs, while spheroids images were acquired at 512×512 pixels, pixel size = 184 nm, dwell time = 2.8 μs. JF_669_, JF_632_, JF_571_ were excited at 1240, 1190, 1040 nm (fixed line) respectively and fluorescence was collected at 650-750 nm, 650-750 nm, 585-750 nm, respectively.

2D HeLa cells as well as spheroids were imaged before and between 20-30 min after hEGF treatment. Cells were segmented using the green channel (EGFP-CAAX). All experiments were performed at ambient room temperature.

### Confocal FLIM microscopy of cysts

Cyst were imaged at 37 °C using the 10× objective (512×512 pixels, pixel size = 328 nm, dwell time = 2.6 μs) exciting at 488 and 669 nm, and collecting at 498-602, 679-750 nm while acquiring FLIM only in the far-red channel. Images were acquired every 2 min for a total of 2 h. Cells were segmented in each image using the green channel (EGFP-CAAX).

### Culture and transfection of primary neuronal co-cultures

The preparation of neuron/glia co-cultures were carried out according to the national guidelines of the Swiss Federal act on animal protection and were approved by the Cantonal Veterinary Office Zurich (license no. ZH229/2024). Pregnant Wistar rats were purchased from ENVIGO, Netherlands. All cell culture media used were from Gibco. The pregnant rat was deeply anesthetized with isoflurane, euthanized by decapitation and embryos were extracted for preparation of primary cultures. Embryos (18 days old) were sacrificed by decapitation and the cerebral cortices were dissected and then washed with 5 mL sterile-filtered PBGA buffer (PBS containing 10 mM glucose, 1 mg mL^−1^ bovine serum albumin and antibiotic-antimycotic 1:100 (10,000 units/ml penicillin; 10,000 µg mL^−1^ streptomycin; 25 µg mL^−1^ amphotericin B). The cortices were cut into small tissue pieces with a sterile scalpel and then digested by incubating in 5 mL sterile filtered papain solution for 15 min at 37 °C. The supernatant was then removed and the tissue was washed twice with complete DMEM/FBS medium (Dulbecco’s modified Eagle’s Medium containing 10% Fetal Bovine Serum and penicillin/streptomycin, 1:100). Then, 3-4 mL of fresh DMEM/FBS was added, and the tissue was carefully and gently triturated and subsequently filtered through a 40 μm cell-strainer. Finally, the neurons were plated at a density of 18,000-22,000 cells per chamber onto the poly D-lysine (Gibco Life Technologies) coated 4 chamber bottom glass dishes (Cellvis) and incubated for 3-4 h at 37°C and 5% CO_2_. Then, the DMEM medium was exchanged with freshly prepared NU-medium (Minimum Essential Medium (MEM) with 15% NU serum, 2% B27 supplement, 15 mM HEPES, 0.45% glucose, 1 mM sodium pyruvate, 2 mM GlutaMAX). The cells were kept in incubator at 37 °C and 5% CO_2_. After 9 days, the cultures were transfected with a total of 1 µg of pAAV-hSyn-WHaloEKAR plasmid DNA or pAAV-hSyn-WHaloEKAR and pAAV-hSyn-mTQ2-Ca-FLITS 500 ng each by the magnetofection method using Lipofectamine 2000 (Invitrogen) and CombiMag (OZ Biosciences) at 37 °C and 5% CO_2_, as described by Buerli et al.^61^

3 days after transfection the neuron co-cultures were labeled with JF_669_-HTL (100 nM for 30 min) at 37 °C in neuron growth medium, washed twice with the same medium, and placed in L15 for imaging.

### Neuron imaging

Neurons were imaged at 37 °C using the 63× objective (512×512 pixels, pixel size = 481 nm, dwell time = 2.8 μs). For single color imaging excitation was performed at 669 nm and fluorescence was collected at 685-820 nm. Dual color imaging was performed under the same conditions with additional excitation at 440 nm and collection from 455-601 nm. 80 MHz repetition rate was used. Images were acquired every 2 min for a total of 30 min and stimulation, or inhibition was performed after the first frame. To determine the fluorescence lifetime of the soma and neurites, individual neurons were segmented using an intensity threshold (overall neuron = soma + neurites). The soma was segmented by hand and the neurites defined as overall neuron - soma.

### Zebrafish experiments

WHaloEKAR-T2A-EGFP mRNA for injection was produced with a mMESSAGE mMACHINE SP6 Transcription Kit (Invitrogen) from a linearized column purified (Qiagen, 74104) DNA fragment. RNA concentration was adjusted to 500 ng µL^−1^ and stored as aliquots at −80 °C.

Zebrafish strains were maintained, grown, and bred according to standard procedures described previously^62^. All experiments were conducted in accordance with the regulations and guidelines of the Veterinary Office of the University of Zürich and the Canton of Zürich, Switzerland, and in compliance with European Union Directive 2010/63/EU. Embryos were staged according to reference^63^. All embryos used in this study were younger than 72 h post-fertilization (hpf), corresponding to a developmental stage prior to sex differentiation; therefore, sex was not considered in the experimental design. One-to two-cell-stage embryos were injected with 0.5 - 1 nL mRNA at a concentration of 250 ng µL^−1^ within 40 min of fertilization. Unfertilized embryos were removed within 12 hpf. At 24 hpf, embryos were screened for sensor expression based on co-expression of EGFP. Embryos were maintained in E3 medium throughout the experiments. To facilitate and accelerate dechorionation, embryos between 24 and 48 hpf were treated with 0.05% pronase for 5 min at RT.

### Labeling of zebrafish embryo, mounting, and imaging

JF_669_-HTL was added at a final concentration of 1 µM from a 1 mM stock to E3 buffer for 2 h. The embryos were then washed for 2 × 2 h with fresh E3. For inhibited condition SU5402 at 1 μM was added 2 h before imaging. For stimulated condition PMA was added to a final concentration of 100 ng mL^−1^ before mounting. Zebrafish embryos were anesthetized using tricaine (0.01%) and embedded in 0.8% low-melting agarose (PeqGOLD Low Melt Agarose, PeqLab Biotechnologie GmbH) on 35 mm imaging glass bottom dishes.

Zebrafish embryos were imaged at 28 °C by confocal FLIM using the 63× objective (512×512 pixels, pixel size = 482 nm, dwell time = 1.8 μs) exciting at 669 nm and collecting at 685-800 nm.

### Software and image processing

Statistical analysis was performed using Python and curves were fit in Prism (GraphPad, version 11.0.0). Images were processed with custom written Python code or ImageJ/Fiji^64,65^.

### Statistics and Reproducibility

Representative microscopy images and time courses are shown. The number of replicates is indicated with the experiments. Replicates stem from distinct samples using independent transfection or stable cells from different passages unless otherwise stated. Details regarding statistical testing are given with the experiments in the Supplementary Figures.

## Data Availability

Plasmids generated in this work will be deposited on Addgene and released upon final publication. The stable cell lines generated are available upon request. Source data are provided with this paper.

## Code Availability

Custom Python scripts to analyze widefield, confocal, and FLIM data will be made available at https://github.com/FreiGroup.

## Acknowledgement

We thank the Frei Research Group for general lab support, assistance with cell culture, and discussion of the manuscript in particular we thank A. Nicoli for providing HaloTag7 protein. We thank L. Lavis and J. Grimm (Janelia Research Campus, HHMI) for providing JF fluorophores. We thank H. Farrants (Janelia Research Campus, HHMI) for discussions on chemigenetic biosensors. We thank S. Kumar and C.-J. Shih (ETH Zurich) for access and assistance to their Quantaurus-QY Absolute PL quantum yield spectrophotometer. We gratefully acknowledge ScopeM for their support & assistance in this work. In particular, we thank Justine Kusche of ScopeM for her support & assistance with 1P and 2P FLIM microscopy. We moreover thank C. Henkel (University of Zurich) for fish care.

We acknowledge funding from the following sources: Swiss National Science Foundation (320030E_224301, 310030L_212508 and 320030-236030 to T.P.); Swiss National Science Foundation (310030_204834 to D.G.); Swiss National Science Foundation (CRSK-3_228999 and CR00I5-235927 to M.S.F.)

## Author contributions

E.D. performed rational engineering; performed *in cellulo* characterization of all WHaloEKAR variants; performed widefield and FLIM microscopy and analysis thereof; performed neuron and zebrafish embryo experiments; R.S.E. purified proteins and performed *in vitro* characterization, performed and analyzed 2P microscopy, generated python scripts for image analysis; D.P. performed multiplexing and cyst experiments and analysis thereof, generated python scripts for image analysis in 3D; Y.Z. and J.J assisted with zebrafish embryo preparation; M.A.B. performed neuron preparation and transfection; T.P. and D.G. supervised work and discussed results; M.S.F. supervised work, generated stable cell lines and spheroids, assisted with FLIM microscopy and analysis thereof; E.D. and M.S.F. conceived and designed the study and wrote the manuscript with input from all authors.

