## Supplementary Information for "A chemigenetic fluorescence lifetime-based biosensor for imaging of ERK activity"

### Table of Content

#### Supplementary Figures

Supplementary Figure S1: Comparison of WHaloEKAR and mNG-ERK-KTR. 2
Supplementary Figure S2: Labeling kinetics of JF669-HTL on WHaloEKAR. 3
Supplementary Figure S3: *In vitro* characterization of WHaloEKAR (left panels) and WHaloEKAR(T7A) (right panels). 4
Supplementary Figure S4: WHaloEKAR labeled with different rhodamine-based fluorophores. 6
Supplementary Figure S5: Characterization of stable cell lines generated. 7
Supplementary Figure S6: Two and three-color multiplexing. 8
Supplementary Figure S7: ERK activity measurements in 3D spheroids. 9
Supplementary Figure S8: ERK activity measurements in MDCK cysts. 10

#### Supplementary Tables

Supplementary Table S1: Biosensor variants generated and tested in this study. 11
Supplementary Table S2: Photophysical properties of purified biosensors. 13
Supplementary Table S3: Temperature dependence on the phase ($\tau_\varphi$) and modulation lifetimes ($\tau_M$ ) of purified biosensors. 14
Supplementary Table S4: Test statistics. 15

#### Supplementary Videos

Supplementary Video S1: WHaloEKAR-$JF_{669}$ in HeLa cells. 16
Supplementary Video S2: mNG-ERK-KTR in HeLa cells. 16
Supplementary Video S3: WHaloEKAR-$JF_{669}$ in mitotic HEK239T cells. 16
Supplementary Video S2: WHaloEKAR-$JF_{669}$ in primary cortical rat neurons. 16

**References** 17

#### Supplementary Figures

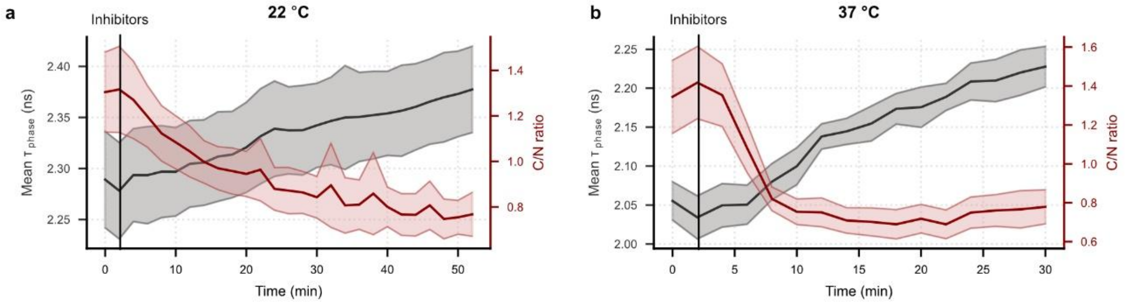

**Supplementary Figure S1:** Comparison of WHaloEKAR and mNG-ERK-KTR. **a, b** Time course measurements of average phase fluorescence lifetime and cytosol-to-nucleus (C/N) ratio upon hEGF stimulation and subsequent inhibition via U0126 and SCH772984 at 22 °C (**a**) and 37 °C (**b**). Both sensors show slower ERK deactivation kinetics at 22 °C compared to 37 °C. mNG-ERK-KTR reaches a plateau faster than WHaloEKAR. This indicates that dephosphorylation of WHaloEKAR is slower than for the KTR, which might be due to the binding of the WW domain to the phospho-site restricting phosphatase access. Time courses are representative of three replicates. Thick solid lines indicate mean responses; shaded areas correspond to 95% confidence interval ($n_{\varphi}$ = 9-10 cells, $n_{KTR}$ = 11-23 cells (**a**); $n_{\varphi}$ = 11-13 cells, $n_{KTR}$ = 11-15 cells (**b**)).

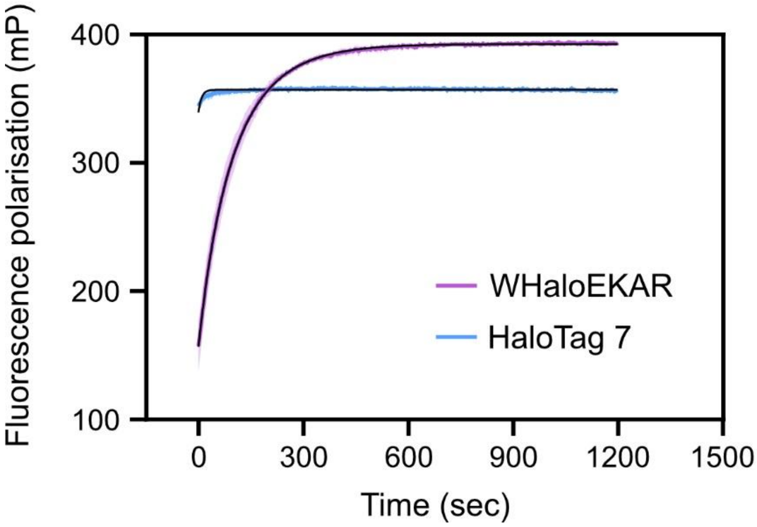

**Supplementary Figure S2:** Labeling kinetics of $JF_{669}$-HTL on WHaloEKAR or HaloTag7. Data are displayed as the means of three technical replicates (purple or blue) and standard deviation (pink or light blue, shaded area), with fitted lines (black). Labeling kinetics with HaloTag7 are not well resolved as $k_{app} > 1 \cdot 10^6\ M^{-1} \cdot s^{-1}$ (Literature $4.03 \pm 0.02\ 10^6\ M^{-1} \cdot s^{-1}$)[1].

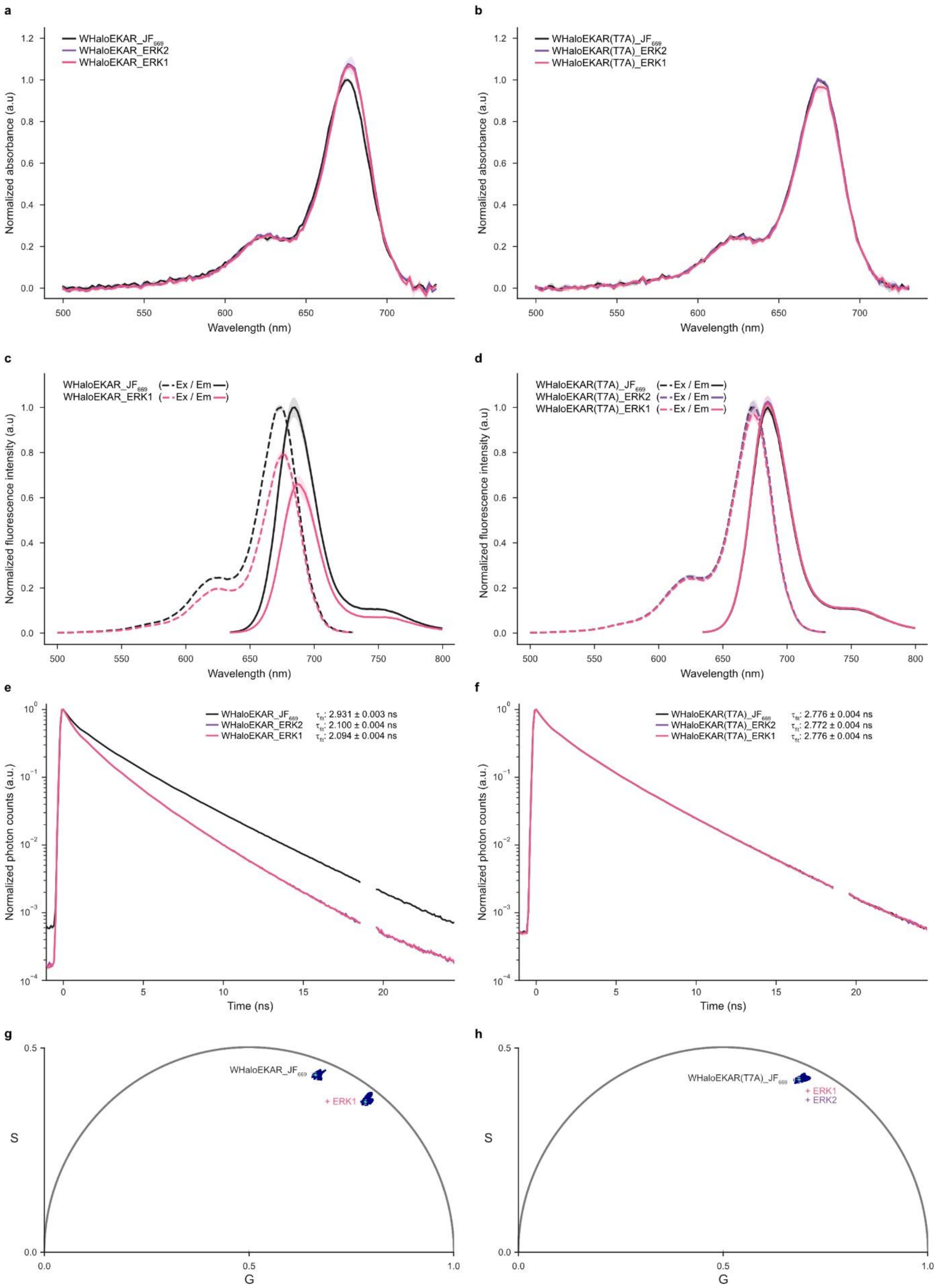

**Supplementary Figure S3**: *In vitro* characterization of WHaloEKAR (left panels) and WHaloEKAR(T7A) (right panels). WHaloEKAR and WHaloEKAR(T7A) were labeled with $JF_{669}$-HTL and data was acquired in the presence or absence of ERK2 or ERK1. Absorbance (**a**, **b**), fluorescence excitation and emission spectra (**c**, **d**), photon counting histogram (**e**, **f**) and phasor plots (**g**, **h**). Absorbance, fluorescence spectra, and photon counting histograms are displayed as the mean and standard deviation from three technical replicates. Phasor plots are displayed from a single replicate. Data is representative of four (WHaloEKAR) and two (WHaloEKAR(T7A)) experiments.

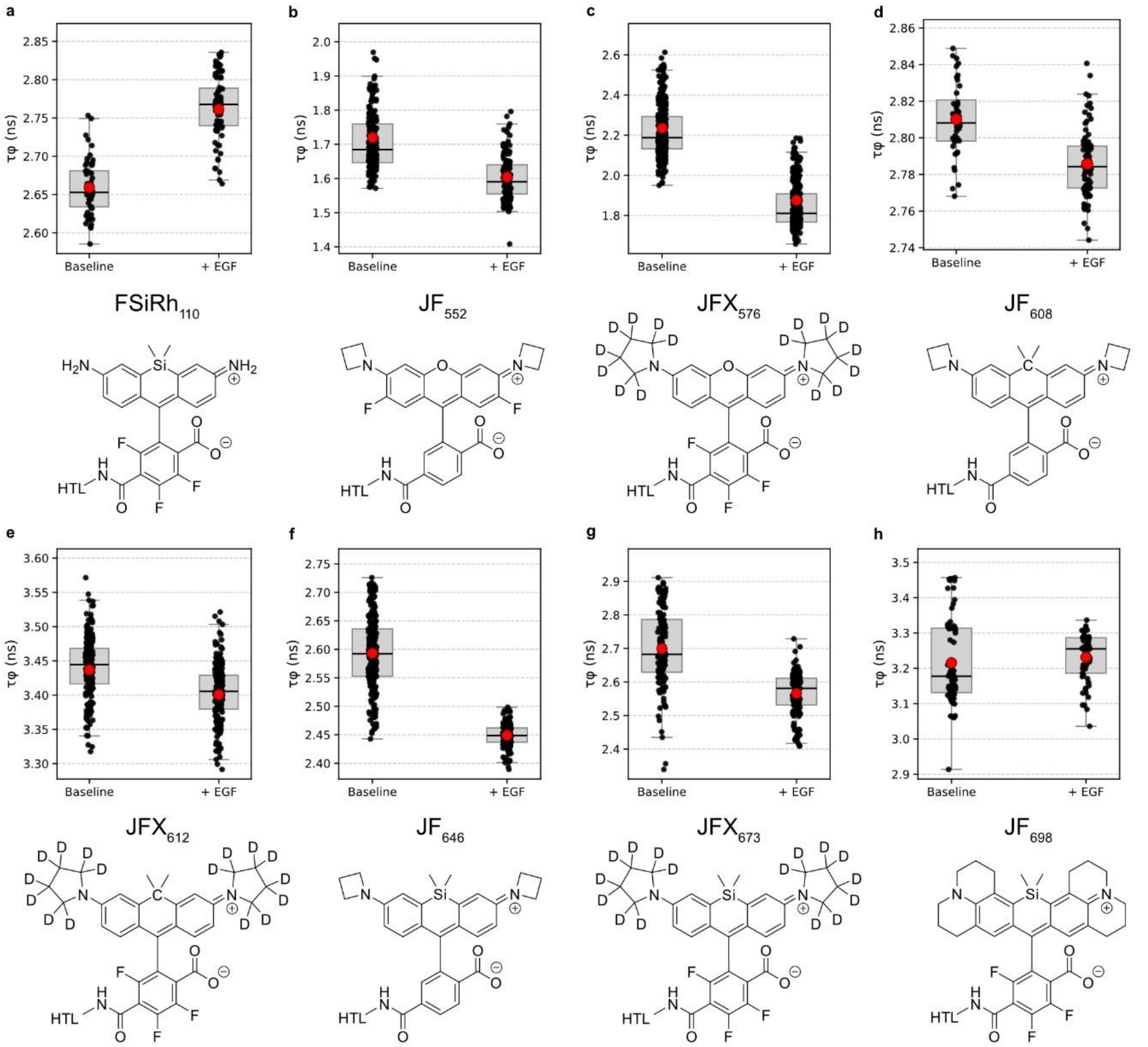

**Supplementary Figure S4:** WHaloEKAR labeled with different rhodamine-based fluorophores. **a-h** Chemical structures and phase fluorescence lifetime of WHaloEKAR labeled with the respective fluorophores before and after hEGF stimulation. Boxplots: Box: 25, 50, 75 percentiles, red dot: mean, whiskers: 1.5 times the interquartile range above and below the 1st and 3rd quartile (*n* = 54 and 83; 203 and 154; 328 and 343; 83 and 40; 193 and 187; 230 and 159; 147 and 143; 57 and 73 cells).

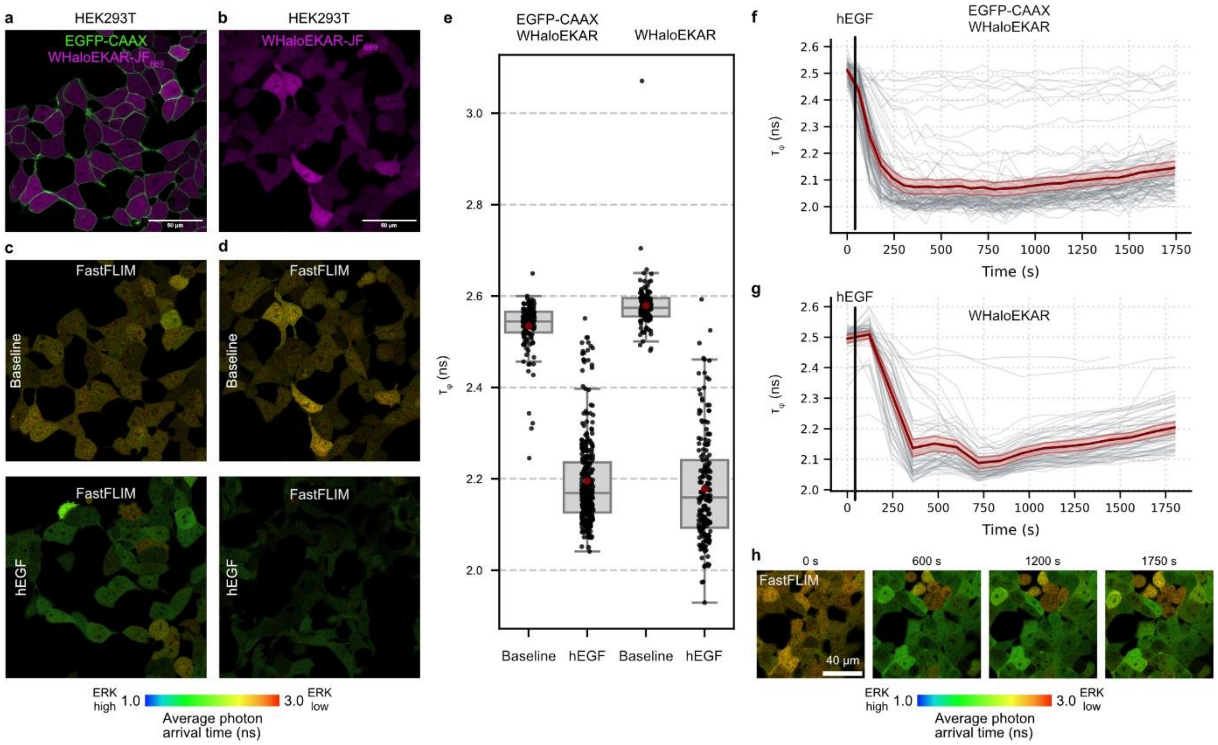

**Supplementary Figure S5:** Characterization of stable cell lines generated. **a-b**, Dual and single channel confocal microscopy images of HEK293T cells stably expressing WHaloEKAR-T2A-EGFP-CAAX (a) or WHaloEKAR labeled with $JF_{669}$-CA. **c-d**, Corresponding FastFLIM images before and after stimulation with hEGF. **e**, Quantification of phase fluorescence lifetime at baseline (30 min starved) and upon hEGF treatment of the two cell lines (*n* = 130, 315, 119, and 187 cells). **f-g**, Average time courses of the stable cell lines upon stimulation with hEGF at the indicated time (*n* = 86 and 56 cells). **h**, FastFLIM images of cells in (f). Unresponsive cells are dividing cells. Boxplots: Box: 25, 50, 75 percentiles, red dot: mean, whiskers: 1.5 times the interquartile range above and below the 1st and 3rd quartile, individual data points are shown on top. Time courses are representative of three replicates, and solid lines indicate addition of drug. Thick solid-red lines indicate mean responses; shaded areas correspond to 95% confidence interval; single-cell curves are shown in gray.

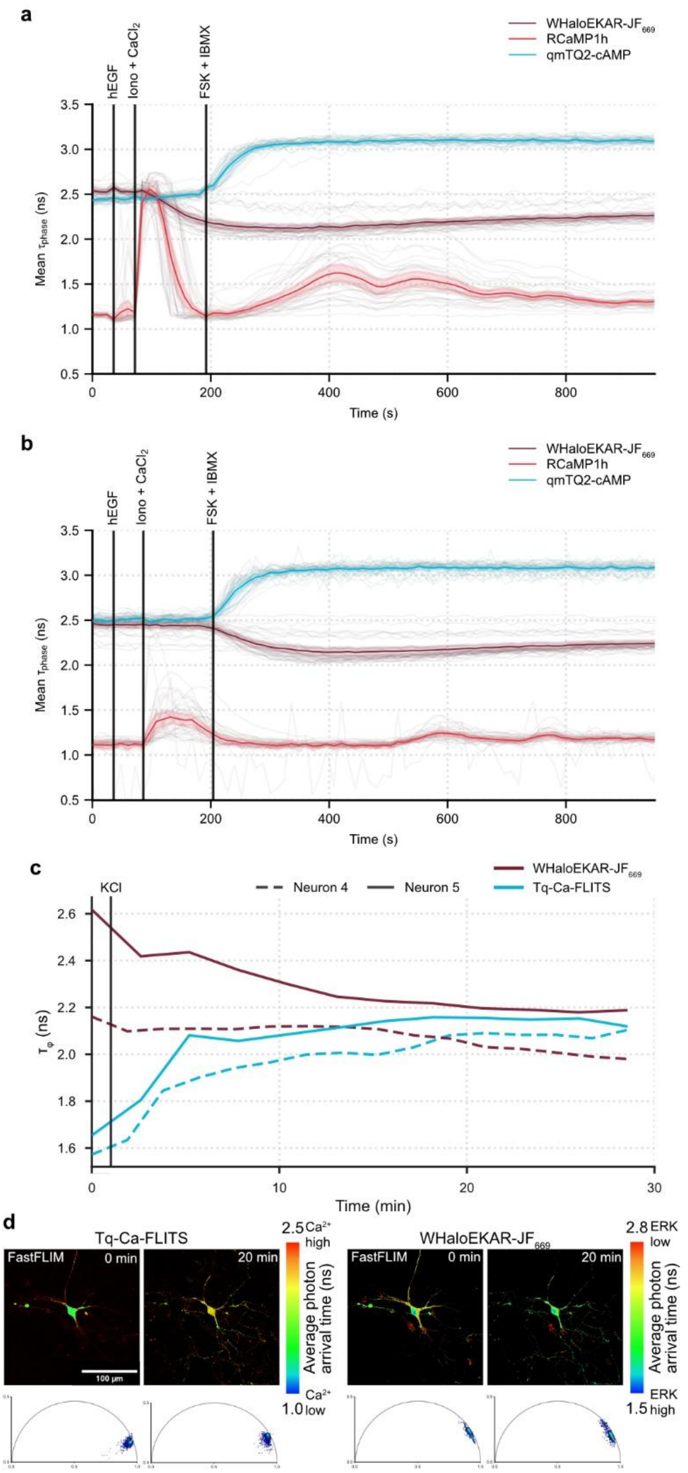

**Supplementary Figure S6:** Two and three-color multiplexing. **a,b** Three-color multiplexing of cAMP (qmTQ2-cAMP, cyan), $Ca^{2+}$ (RCaMP1h, red) and ERK activity (WHaloEKAR-$JF_{669}$, dark-red) in HEK239T cells stimulated with hEGF, ionomycin + $CaCl_2$ (1 µM, 5 mM), and FSK + IBMX (50 µM/ 100 µM). All three activities are read out by FLIM (*n* = 41 (a), 45 (b) cells). **c,** Two-color multiplexing of $Ca^{2+}$ (Tq-Ca-FLITS, cyan) and ERK activity (WHaloEKAR-$JF_{669}$, dark-red) in primary cortical neurons stimulated with KCl (50 µM). Both activities are read out by FLIM (*n* = 2 cells). **d,** Representative FastFLIM images and phasor plots of neurons imaged in Figure 5c and S6c. a, b. Time courses are representative of three replicates. Thick solid lines indicate mean responses; shaded areas correspond to 95% confidence interval; single-cell curves are shown in gray. c, Single curves are shown for individual neurons.

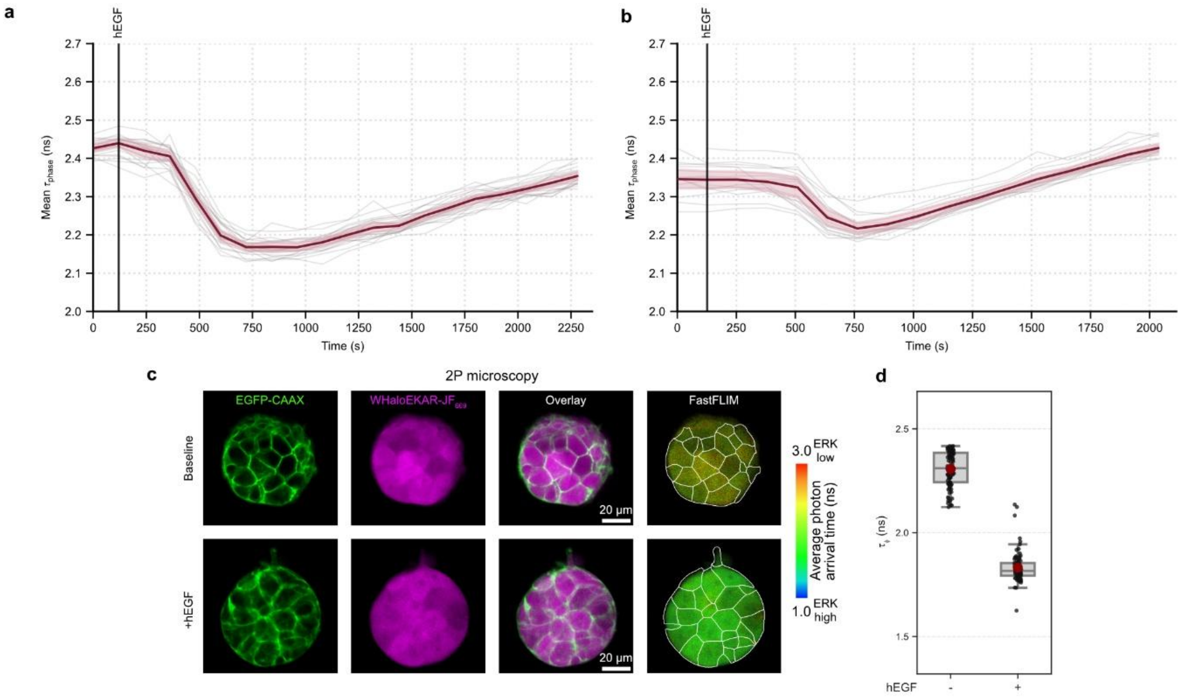

**Supplementary Figure S7:** ERK activity measurements in 3D spheroids. **a, b** Representative time-course measurements of phase fluorescence lifetime of individual HEK293T cells in 4-day old spheroids. Stimulation with hEGF leads to a transient increase in ERK activity (*n* = 20 (**a**), 15 (**b**) cells). **c**, Representative confocal 2P intensity and FastFLIM images of the median plane 4-day HEK293T spheroids co-expressing WHaloEKAR and EGFP-CAAX labeled with $JF_{669}$-HTL, before and after hEGF stimulation. Individual color channels as well as overlay are given. Outline of cells segmented are overlaid over the FastFLIM image. Scale bars, 20 µm. **d**, Quantification of phase fluorescence lifetime measured by 2P FLIM at baseline (30 min starved) and upon hEGF treatment (*n* = 89 and 82 cells). Time courses are representative of three replicates. Thick solid lines indicate mean responses; shaded areas correspond to 95% confidence interval; single-cell curves are shown. Boxplots: Box: 25, 50, 75 percentiles, red dot: mean, whiskers: 1.5 times the interquartile range above and below the $1^{st}$ and $3^{rd}$ quartile, individual data points are shown on top. Data are pooled from two experiments containing 2-3 spheroids per experiment

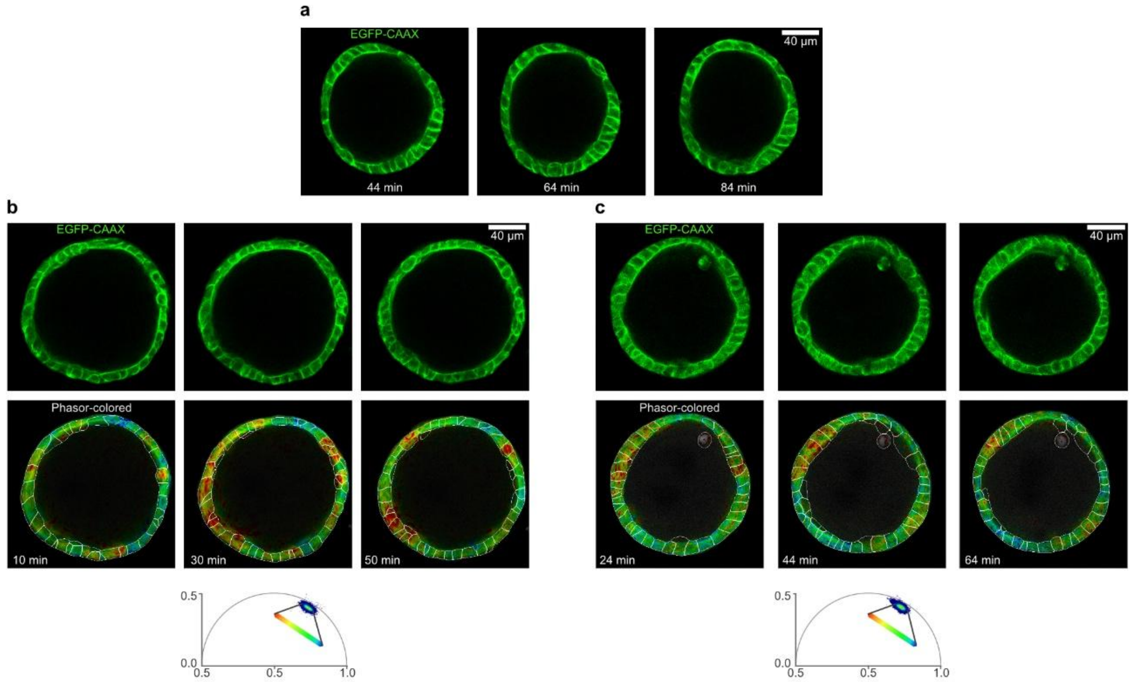

**Supplementary Figure S8:** ERK activity measurements in MDCK cysts. **a** GFP channel of MDCK cyst shown in Figure 6d. This channel was used to segment individual cells. Scale bar, 40 µm. **b-c**, Representative EGFP signal and phasor-colored images of the far-red channel of 6-day old MDCK cyst stably expressing WHaloEKAR and EGFP-CAAX labeled with $JF_{669}$-HTL. FLIM was performed for 2 h. Selected time points are shown. Phasor used to define the look up table is given below. Outline of individual cells as defined by the EGFP channel are given. Scale bars, 40 µm.

#### Supplementary Tables

**Supplementary Table S1:** Biosensor variants generated and tested in this study. The best variant in each generation is highlighted in bold. cpHT refers to cpHaloTag[141-147]. Variant 5.1 was dubbed WHaloEKAR.

| # | Domain1 | L1 | HaloTag-Mutations | L2 | Domain2 |
|---|---|---|---|---|---|
| *1.1* | ERK Substrate | QMR | cpHT-Q165W | TIS | WW-Domain |
| *1.2* | ERK Substrate | QMR | cpHT-P174W | TIS | WW-Domain |
| *1.3* | ERK Substrate | QMR | cpHT-M175W | TIS | WW-Domain |
| *1.4* | ERK Substrate | QMR | cpHT-G171W | TIS | WW-Domain |
| *1.5* | ERK Substrate | MRR | cpHT-Q165W | TIS | WW-Domain |
| *1.6* | ERK Substrate | MRR | cpHT-P174W | TIS | WW-Domain |
| *1.7* | ERK Substrate | MRR | cpHT-M175W | TIS | WW-Domain |
| *1.8* | ERK Substrate | MRR | cpHT-G171W | TIS | WW-Domain |
| *1.9* | ERK Substrate | WRR | cpHT-Q165W | TIS | WW-Domain |
| *1.10* | ERK Substrate | WRR | cpHT-P174W | TIS | WW-Domain |
| *1.11* | ERK Substrate | WRR | cpHT-M175W | TIS | WW-Domain |
| *1.12* | ERK Substrate | WRR | cpHT-G171W | TIS | WW-Domain |
| *1.13* | WW-Domain | QMR | cpHT-Q165W | TIS | ERK Substrate |
| *1.14* | WW-Domain | QMR | cpHT-P174W | TIS | ERK Substrate |
| *1.15* | WW-Domain | QMR | cpHT-M175W | TIS | ERK Substrate |
| *1.16* | WW-Domain | QMR | cpHT-G171W | TIS | ERK Substrate |
| *1.17* | WW-Domain | MRR | cpHT-Q165W | TIS | ERK Substrate |
| *1.18* | WW-Domain | MRR | cpHT-P174W | TIS | ERK Substrate |
| *1.19* | WW-Domain | MRR | cpHT-M175W | TIS | ERK Substrate |
| *1.20* | WW-Domain | MRR | cpHT-G171W | TIS | ERK Substrate |
| ***1.21*** | **WW-Domain** | **WRR** | **cpHT-Q165W** | **TIS** | **ERK Substrate** |
| *1.22* | WW-Domain | WRR | cpHT-P174W | TIS | ERK Substrate |
| *1.23* | WW-Domain | WRR | cpHT-M175W | TIS | ERK Substrate |
| *1.24* | WW-Domain | WRR | cpHT-G171W | TIS | ERK Substrate |
| *2.1* | WW-Domain | ARR | cpHT-Q165W | TIS | ERK Substrate |
| *2.2* | WW-Domain | WAR | cpHT-Q165W | TIS | ERK Substrate |
| *2.3* | WW-Domain | WRA | cpHT-Q165W | TIS | ERK Substrate |
| *2.4* | WW-Domain | AWRR | cpHT-Q165W | TIS | ERK Substrate |
| *2.5* | WW-Domain | WRRA | cpHT-Q165W | TIS | ERK Substrate |
| *2.6* | WW-Domain | AWRRA | cpHT-Q165W | TIS | ERK Substrate |
| *2.7* | WW-Domain | RR | cpHT-Q165W | TIS | ERK Substrate |
| *2.8* | WW-Domain | WRR | cpHT-Q165W-Δ147 | TIS | ERK Substrate |
| *2.9* | WW-Domain | WRR | cpHT-Q165W-Δ147-148 | TIS | ERK Substrate |
| ***2.10*** | **WW-Domain** | **WRR** | **cpHT-Q165W** | **AIS** | **ERK Substrate** |
| *2.11* | WW-Domain | WRR | cpHT-Q165W | TAS | ERK Substrate |
| *2.12* | WW-Domain | WRR | cpHT-Q165W | TIA | ERK Substrate |
| *2.13* | WW-Domain | WRR | cpHT-Q165W | ATIS | ERK Substrate |
| *2.14* | WW-Domain | WRR | cpHT-Q165W | TISA | ERK Substrate |
| *2.15* | WW-Domain | WRR | cpHT-Q165W | ATISA | ERK Substrate |

| | | | | | |
|---|---|---|---|---|---|
| *2.16* | WW-Domain | WRR | cpHT-Q165W | IS | ERK Substrate |
| *2.17* | WW-Domain | WRR | cpHT-Q165W-Δ141 | TIS | ERK Substrate |
| ***2.18*** | **WW-Domain** | **WRR** | **cpHT-Q165W-Δ140-141** | **TIS** | **ERK Substrate** |
| ***3.1*** | **WW-Domain** | **WRA** | **cpHT-Q165W** | **AIS** | **ERK Substrate** |
| *3.2* | WW-Domain | WRA | cpHT-Q165W | TIA | ERK Substrate |
| *3.3* | WW-Domain | WRA | cpHT-Q165W-Δ147-148 | TIS | ERK Substrate |
| *3.4* | WW-Domain | AWRR | cpHT-Q165W | AIS | ERK Substrate |
| *3.5* | WW-Domain | AWRR | cpHT-Q165W | TIA | ERK Substrate |
| *3.6* | WW-Domain | AWRR | cpHT-Q165W-Δ147-148 | TIS | ERK Substrate |
| *4.1* | WW-Domain-Y23F | WRA | cpHT-Q165W | AIS | ERK Substrate |
| *4.2* | WW-Domain-W34F | WRA | cpHT-Q165W | AIS | ERK Substrate |
| ***4.3*** | **WW-Domain** | WRA | **cpHT-M86L-Q165W** | AIS | **ERK Substrate** |
| *4.4* | WW-Domain-Y23F-W34F | WRA | cpHT-Q165W | AIS | ERK Substrate |
| *4.5* | WW-Domain-W34F | WRA | cpHT-M86L-Q165W | AIS | ERK Substrate |
| *4.6* | WW-Domain-Y23F-W34F | WRA | cpHT-M86L-Q165W | AIS | ERK Substrate |
| ***5.1*** | **WW-Domain-Δ54** | **WRA** | **cpHT-M86L-Q165W** | **AIS** | **ERK Substrate** |
| *5.2* | WW-Domain | WRAA | cpHT-M86L-Q165W | AIS | ERK Substrate |
| *5.3* | WW-Domain | WRAAA | cpHT-M86L-Q165W | AIS | ERK Substrate |
| *5.4* | WW-Domain | WRA | cpHT-M86L-Q165W-Δ147-148 | AIS | ERK Substrate |
| *5.5* | WW-Domain | WRA | cpHT-M86L-Q165W-Δ147 | AIS | ERK Substrate |
| *5.6* | WW-Domain | WRA | cpHT-M86L-Q165W | AAIS | ERK Substrate |
| *5.7* | WW-Domain-Δ54 | WRA | cpHT-M86L-Q165W-Δ147-148 | AIS | ERK Substrate |
| *5.8* | WW-Domain-Δ54 | WRA | cpHT-M86L-Q165W-Δ147 | AIS | ERK Substrate |
| *5.9* | WW-Domain-Δ54 | WRA | cpHT-M86L-Q165W | AAIS | ERK Substrate |
| *5.10* | WW-Domain | WRAA | cpHT-M86L-Q165W-Δ147-148 | AIS | ERK Substrate |
| *5.11* | WW-Domain | WRAA | cpHT-M86L-Q165W-Δ147 | AIS | ERK Substrate |
| *5.12* | WW-Domain | WRAA | cpHT-M86L-Q165W | AAIS | ERK Substrate |
| *5.13* | WW-Domain | WRAAA | cpHT-M86L-Q165W-Δ147-148 | AIS | ERK Substrate |
| *5.14* | WW-Domain | WRAAA | cpHT-M86L-Q165W-Δ147 | AIS | ERK Substrate |
| *5.15* | WW-Domain | WRAAA | cpHT-M86L-Q165W | AAIS | ERK Substrate |

**Supplementary Table S2:** Photophysical properties of purified biosensors. WHaloEKAR and WHaloEKAR(T7A). They were labeled with $JF_{669}$-HTL in the apo state and phosphorylated with ERK1 or ERK2. Absorbance, excitation, and emission maxima are given along with the extinction coefficient at 678 nm (ε), the quantum yield (φ), the phase ($\tau_\varphi$) and modulation lifetime ($\tau_M$), their standard deviations (SD) and the lifetime change compared to the apo state ($\Delta\tau_\varphi$ and $\Delta\tau_M$). Data for WHaloEKAR stems from 4 experimental replicates and for WHaloEKAR(T7A) from 2 experimental replicates. All data was measured at room temperature.

| | *Abs* *nm* | *Ex* *nm* | *Em* *nm* | *ε* $M^{-1}\cdot cm^{-1}$ | *φ* | *Brightness* $M^{-1}\cdot cm^{-1}$ | $\tau_\varphi$ *ns* | $\tau_M$ *ns* | $SD\tau_\varphi$ *ns* | $SD\tau_M$ *ns* | $\Delta\tau_\varphi$ *ns* | $\Delta\tau_M$ *ns* |
|---|---|---|---|---|---|---|---|---|---|---|---|---|
| *WHaloEKAR (Apo)* | 675 | 671 | 686 | 103178 | 0.22 | 22.3 | 2.71 | 3.21 | 0.06 | 0.07 | N/A | N/A |
| *WHaloEKAR (ERK2)* | 677 | 673 | 689 | 106806 | 0.14 | 15.0 | 1.91 | 2.37 | 0.06 | 0.10 | 0.80 | 0.84 |
| *WHaloEKAR (ERK1)* | 677 | 673 | 689 | 114794 | 0.14 | 16.0 | 1.90 | 2.36 | 0.06 | 0.10 | 0.81 | 0.85 |
| *WHaloEKAR(T7A) (Apo)* | 675 | 672 | 687 | 100515 | 0.20 | 19.8 | 2.53 | 3.05 | 0.08 | 0.10 | N/A | N/A |
| *WHaloEKAR(T7A) (ERK2)* | 675 | 673 | 687 | 97499 | 0.20 | 19.7 | 2.53 | 3.05 | 0.09 | 0.11 | 0.00 | 0.00 |
| *WHaloEKAR(T7A) (ERK1)* | 675 | 672 | 687 | 98348 | 0.20 | 20.0 | 2.54 | 3.05 | 0.09 | 0.11 | 0.01 | 0.01 |

**Supplementary Table S3:** Temperature dependence on the phase ($\tau_\varphi$) and modulation lifetimes ($\tau_M$) of purified biosensors. Values are given for 22 °C, 28 °C and 37 °C. Samples were labeled with $JF_{669}$-HTL in the apo state and phosphorylated with ERK1 or ERK2.

| | *Temperature (°C)* | *$\tau_\varphi$ ns* | *$\tau_M$ ns* | *$SD\tau_\varphi$ ns* | *$SD\tau_M$ ns* | *$\Delta\tau_\varphi$ ns* | *$\Delta\tau_M$ ns* |
|---|---|---|---|---|---|---|---|
| *WHaloEKAR (Apo)* | 22 | 2.67 | 3.17 | 0.06 | 0.05 | N/A | N/A |
| | 28 | 2.64 | 3.13 | 0.04 | 0.05 | N/A | N/A |
| | 37 | 2.46 | 2.98 | 0.04 | 0.05 | N/A | N/A |
| *WHaloEKAR (ERK2)* | 22 | 1.93 | 2.37 | 0.03 | 0.04 | 0.74 | 0.79 |
| | 28 | 1.87 | 2.33 | 0.03 | 0.05 | 0.77 | 0.81 |
| | 37 | 1.79 | 2.25 | 0.03 | 0.05 | 0.67 | 0.73 |
| *WHaloEKAR (ERK1)* | 22 | 1.93 | 2.37 | 0.03 | 0.04 | 0.74 | 0.80 |
| | 28 | 1.87 | 2.33 | 0.03 | 0.04 | 0.77 | 0.81 |
| | 37 | 1.79 | 2.25 | 0.04 | 0.05 | 0.67 | 0.73 |
| *WHaloEKAR(T7A) (Apo)* | 22 | 2.52 | 3.02 | 0.04 | 0.04 | N/A | N/A |
| | 28 | 2.45 | 2.97 | 0.04 | 0.05 | N/A | N/A |
| | 37 | 2.30 | 2.82 | 0.04 | 0.05 | N/A | N/A |
| *WHaloEKAR(T7A) (ERK2)* | 22 | 2.52 | 3.02 | 0.03 | 0.04 | 0.00 | 0.00 |
| | 28 | 2.46 | 2.97 | 0.04 | 0.05 | -0.01 | -0.01 |
| | 37 | 2.30 | 2.83 | 0.04 | 0.05 | -0.01 | -0.01 |
| *WHaloEKAR(T7A) (ERK1)* | 22 | 2.52 | 3.02 | 0.04 | 0.04 | 0.00 | 0.00 |
| | 28 | 2.46 | 2.97 | 0.04 | 0.05 | -0.01 | -0.01 |
| | 37 | 2.31 | 2.83 | 0.04 | 0.05 | -0.01 | -0.01 |

**Supplementary Table S4:** Test statistics. Welch's one-way ANOVA (two-sided, $F$ = 97.7, DFn = 14, DFd = 69.8) comparing fluorescence lifetime at different subcellular locations and stimulation conditions. Data for relevant individual comparisons are given.

| *Group 1* | *Group 2* | *Mean Diff (G1 - G2)* | *Welch t* | *Welch df* | *Welch p-val (Bonferroni)* |
|---|---|---|---|---|---|
| *FYVE_Bas* | NES_Bas | -0.015 | 44.3 | 0.659497 | 1 |
| *LAMP1_Bas* | NES_Bas | -0.282 | 19.6 | 3.38E-05 | 0.0035 |
| *Lyn_Bas* | NES_Bas | -0.473 | 22.8 | 2.14E-08 | 2.25E-06 |
| *NES_Bas* | TOMM_Bas | 0.364 | 12.1 | 3.91E-05 | 0.0041 |
| *FYVE_Bas* | FYVE_EGF | 0.428 | | | 8.26E-11 |
| *LAMP1_Bas* | LAMP1_EGF | 0.221 | 26.6 | 7.87E-13 | 0.0461 |
| *Lyn_Bas* | Lyn_EGF | 0.178 | 14.8 | 0.000439 | 0.5349 |
| *NES_Bas* | NES_EGF | 0.395 | 21.4 | 0.005094 | 3.67E-13 |
| *TOMM_Bas* | TOMM_EGF | -0.095 | 32.1 | 3.49E-15 | 1 |
| | | | 26.9 | 0.256198 | |
| *FYVE_Bas* | FYVE_inhibit | -0.092 | | | 0.2617 |
| *LAMP1_Bas* | LAMP1_inhibit | -0.076 | 34.8 | 0.002492 | 1 |
| *Lyn_Bas* | Lyn_inhibit | -0.008 | 15.3 | 0.145025 | 1 |
| *NES_Bas* | NES_inhibit | -0.057 | 20.0 | 0.880344 | 1 |
| *TOMM_inhibit* | TOMM_Bas | -0.079 | 44.3 | 0.086884 | 1 |

#### Supplementary Videos

**Supplementary Video S1:** WHaloEKAR-$JF_{669}$ in HeLa cells. FastFLIM video of HeLa cells expressing WHaloEKAR-T2A-mNG-ERK-KTR labeled with $JF_{669}$-HTL over 38 min. Stimulation with hEGF was performed at 2 min. Average photon arrival times are scaled between 1 and 3 ns. Scale bar, 50 µm.

**Supplementary Video S2:** mNG-ERK-KTR in HeLa cells. Confocal intensity video of the green channel of HeLa cells expressing WHaloEKAR-T2A-mNG-ERK-KTR labeled with $JF_{669}$-HTL over 38 min. Stimulation with hEGF was performed at 2 min. Average photon arrival times are scaled between 1 and 3 ns. Scale bar, 50 µm.

**Supplementary Video S3:** WHaloEKAR-$JF_{669}$ in mitotic HEK239T cells. FastFLIM video of HEK293T cells stably expressing WHaloEKAR labeled with $JF_{669}$-HTL over 28 min. Stimulation with hEGF was performed at 2 min. Average photon arrival times are scaled between 1 and 3 ns. Scale bar, 20 µm.

**Supplementary Video S4**: WHaloEKAR-$JF_{669}$ in primary cortical rat neurons. FastFLIM video of a primary cortical rat neuron expressing WHaloEKAR labeled with $JF_{669}$-HTL over 58 min. Stimulation with BDNF was performed after the first frame and inhibition with U0126 and SCH772984 after 28 min. Average photon arrival times are scaled between 1 and 3 ns. Scale bar, 50 µm.
