## Supplementary figures and images for "A chemigenetic fluorescence lifetime-based biosensor for imaging of ERK activity"

### Video S3

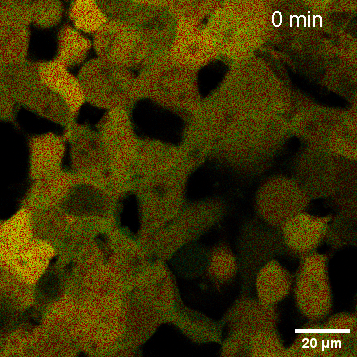
